# Identification of Mirror Repeats in Viral Genomes using FPCB Analysis

**DOI:** 10.1101/2023.04.13.536685

**Authors:** Pooja Yadav, Jyoti Kumari, Priyanka Yadav, Rachna Yadav, Shivani Yadav, Dinesh Sharma, Amrita Singh, Barkha Sehrawat, Manisha Yadav, Sandeep Yadav

## Abstract

The majority of living domains consist of DNA as genetic material with the minor exception of viruses. The unique nature of every species determines by its unique pattern of genome or gene products. The genomic features become an evident example of evolutionary study also. Different types of repeat patterns are observed in genomes of living domains including human beings whose two third portion of the genome is repetitive. Among the varied type of repeat sequences Mirror Repeats (MR) play crucial roles at the genetic level in every species. The major focus of our research is on identification & to check the distribution of mirror repeat. For this, we employed a bioinformatics-based approach refer as FASTA PARALLEL COMPLEMENT BLAST (FPCB) to identify unique mirror repeat (MR) sequences in some selected viral genomes from three different categories (Animal, Plant & Human). The identified repeats vary in their length as well as found to be distributed throughout the selected viral genomes. The maximum no of MR were reported in the case of Dengue virus (229) & minimum is in the case of TMV (97). In the remaining selected viruses - HCV, HPV, HTLV-1, PVY, Rabies virus 178, 156, 175, 203 & 204 MR sequences were reported. These sequences can be utilized in many ways like in molecular diagnosis, drug delivery target as well as evolutionary study, etc. The present research also helps in the development of novel tools of bioinformatics to study mirror repeats and their functional perspective in the context of their occurrence in all domains.

## Introduction

The regulatory molecule (DNA) is responsible for genetic diversity among species from viruses to the complex living system including human beings [1,2]. The genetics of an organism can be depending upon various factors including environmental ones as well as at the level of genes or genome. The arrangements of base pair patterns in the genes or genomes of every domain resulting the diversification of species or strains [3,4]. These unique patterns form various types of repeats. Every type performs different kinds of functions at the molecular level & contributes to normal cellular functions. The most common types include inverted repeats, tandem repeats, satellite DNA, SNP’s, transposable elements and palindrome sequences, etc [5,6,7] (See **Figure 1**). These repeats were found to be associated with varied functions like genomic structural determination, gene expression regulation, binding sites for many enzymes, heterochromatin region formation & genomic evolution, and associated with many human diseases [8-11]. Among the above-mentioned repeats a unique pattern refers to as Mirror repeat (MR) is also being reported over time [12-14]. A Mirror repeat is defined as when one part of its sequence share homology with the rest part of the same sequence. For example ATGCGGGGCGTA, in this sequence its one part ATGCGG share homology with another part. Mirror repeats have a center of symmetry on the same strand of the genome. These sequences (MR’s) were reported to be associated with various kinds of functions in the genome. One of the major contributions of these repeats is in the formation of triplex DNA & unusual genomic structures by folding the genome [16,17]. They are found to be associated with genetic disorders [18, 19] & also reported to be responsible for genomic instability [20]. Mirror repeats form the Non-B forms of the DNA by non-Watson Crick base pairing. The present investigation focuses on the identification of Mirror repeats using a bioinformatics - based approach. The approach refers to as FASTA PARALLEL COMPLEMENT BLAST (FPCB) which follows some simple steps to identify MR’s [21]. We select viral genomes from the plant (TMV & Potato Virus Y), human (HPV, HCV & HTLV-1) & animal categories (Dengue virus & Rabies virus) because the size of viral genomes is small and easily handled during manual bioinformatics analysis. The selected viral genomes from different categories were undergone FPCB analysis in which we try to identify mirror repeats & check their distribution in the genomes. This study will be helpful for more insights on MR sequences in the context of their functions, roles as well as the pattern of occurrence in every species.

**Figure 1.**
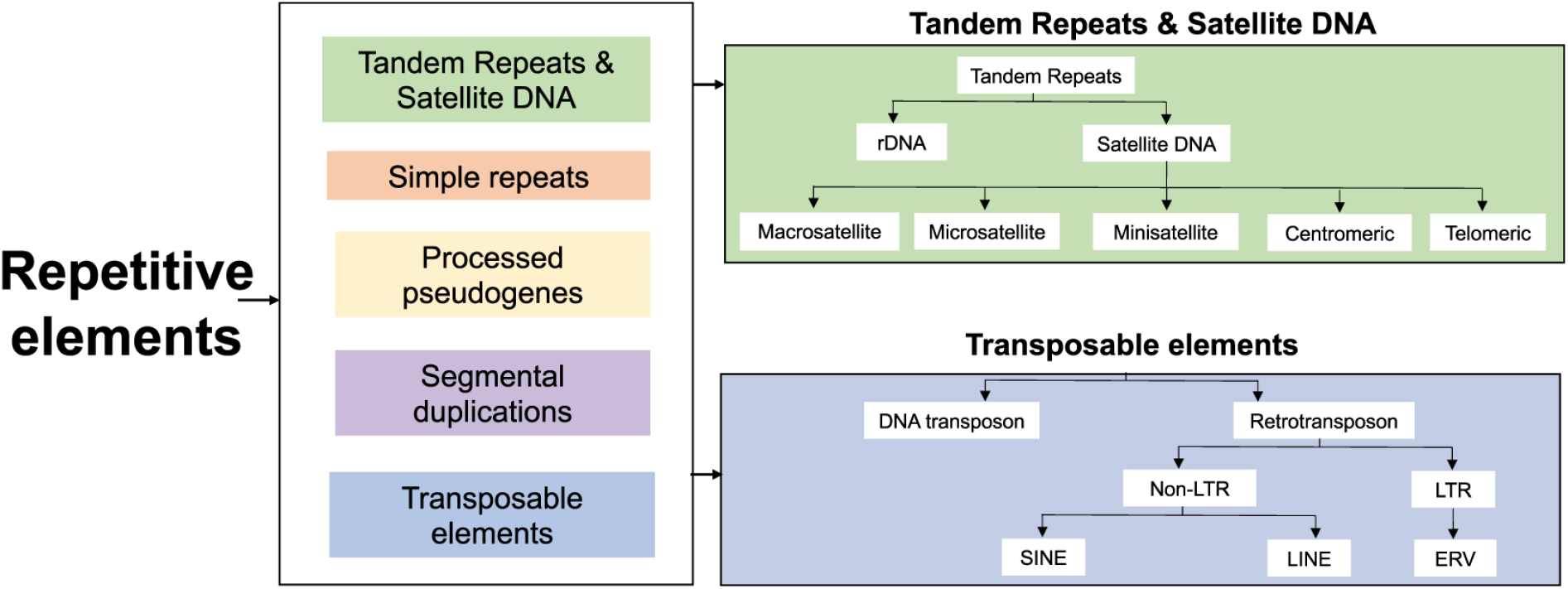
Represent the various types of repeat elements. (Adapted from Planken, A. *et al*. 2019 [15])

## Material & Methods

To identify the Mirror repeat sequences a four-step bioinformatics-based approach is utilized. The approach refers to as FPCB (FASTA PARALLEL COMPLEMENT BLAST) analysis. In its first step the genome sequence of the selected virus is downloaded from NCBI [22], followed by the division of the same into 1000 base pairs regions to ensure the maximum identification of MR sequences with minimum length. In a further step, the divided regions act as a query sequence whose complement sequence retrieve from the Reverse complement tool, which acts as our subject sequence. The final step includes the BLAST analysis between Query & Subject sequences. If the position no of the query & subject sequence is exactly reverse then it will be a Mirror repeat [21]. The flow model of the methodology is given below in **Figure 2**[23]-

**Figure 2.**
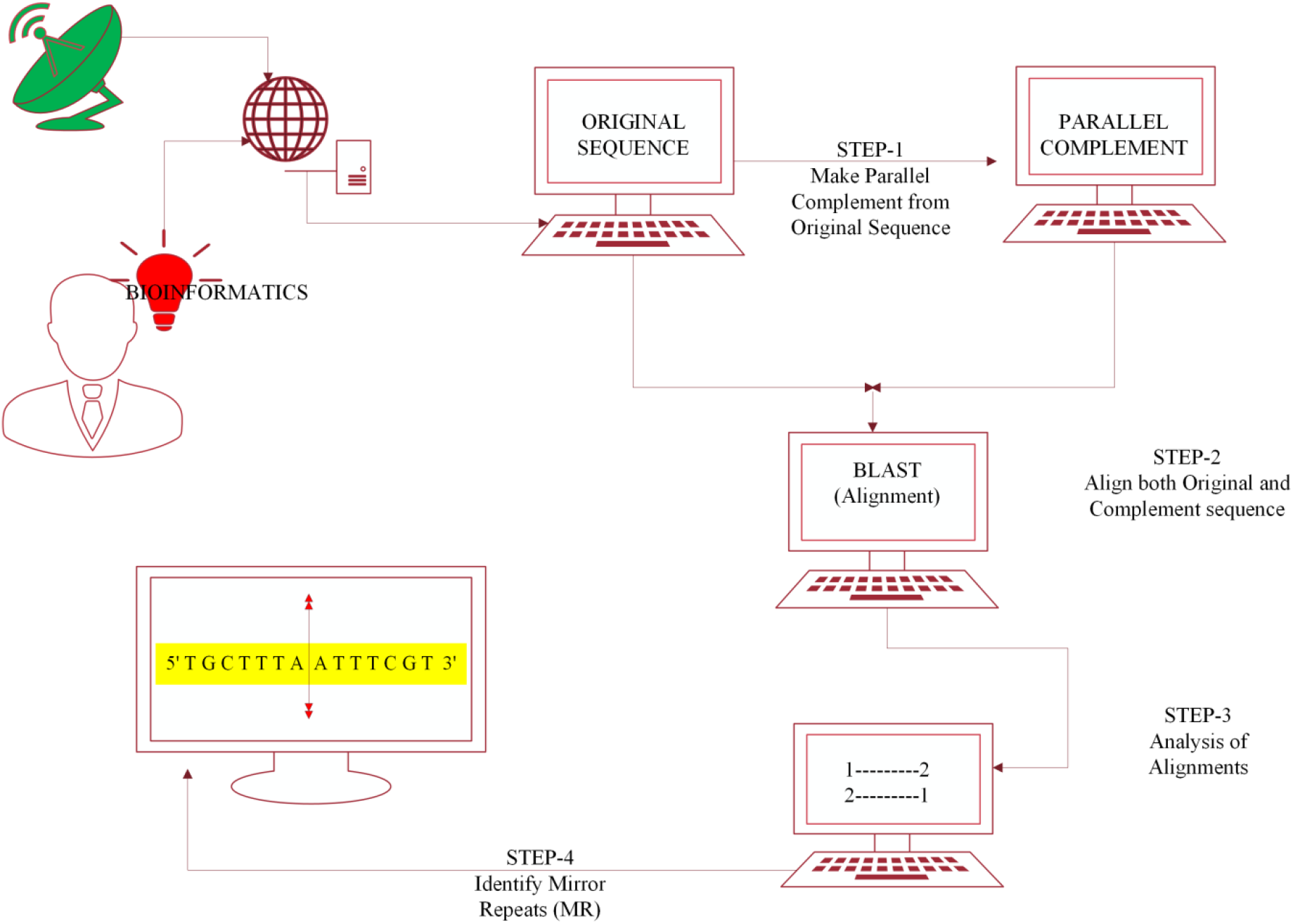
Depicted the steps involve in FPCB in which the original sequence retrieved from NCBI followed by making its complement sequence & BLAST analysis between original & complement sequence. The exact reverse position no of the original & complement sequence represents an MR sequence. (Adapted from Yadav, S *et al* 2022 [23])

**Figure 3.**
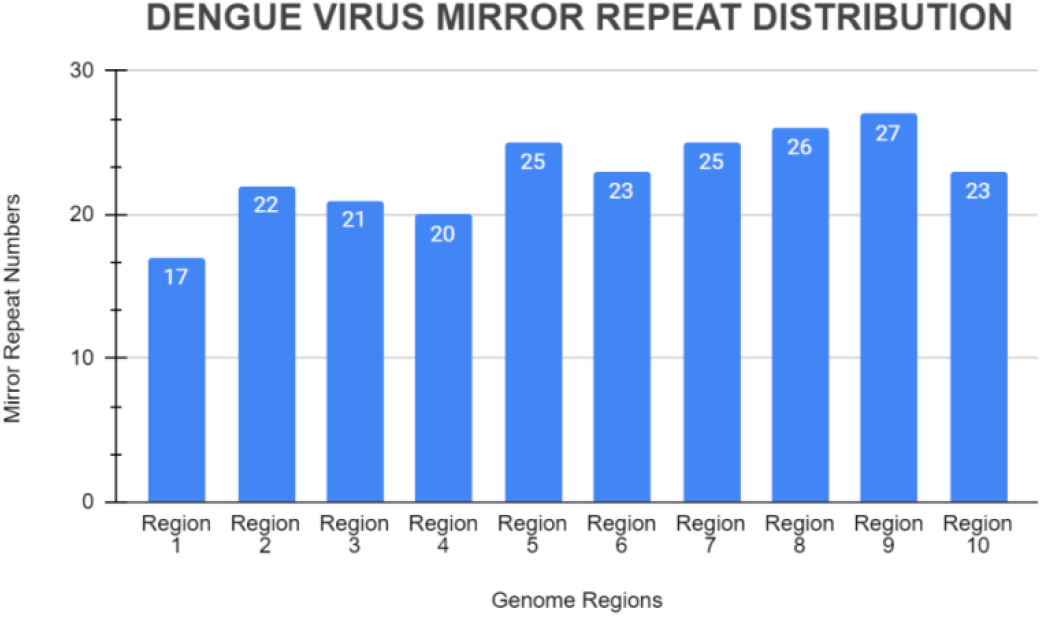
Represent the MR distribution in Dengue virus. The maximum no of MR sequences was reported in Region 9 (27) while the minimum no is in Region 4 (20).

**Figure 4.**
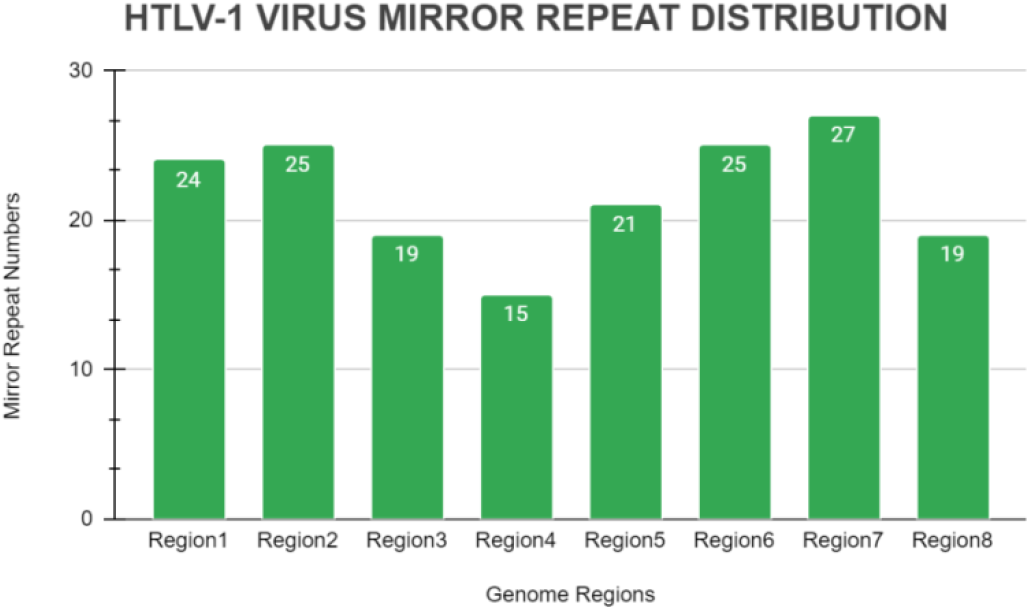
Represent the MR distribution in HTLV-1. The maximum no of MR sequences was found in Region 7 (27) while the minimum no is in Region 4 (15).

**Figure 5.**
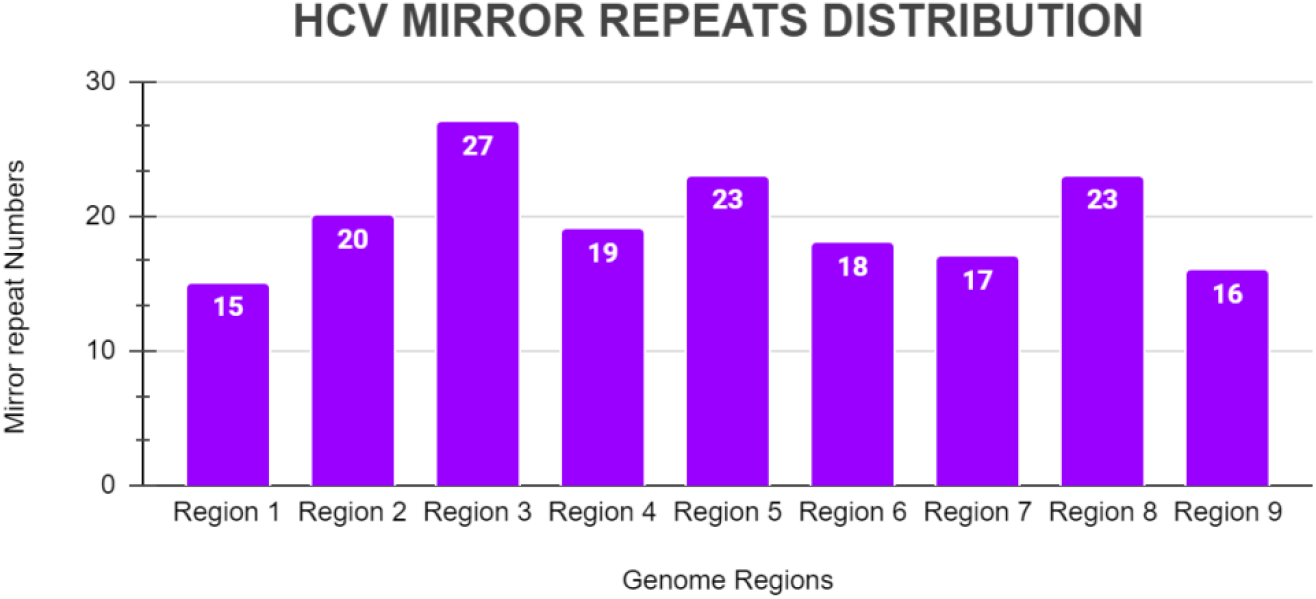
Represent the MR distribution in HCV. The maximum & minimum no of MR sequences were reported in Region 3 (27) & Region 1 (15) respectively.

**Figure 6.**
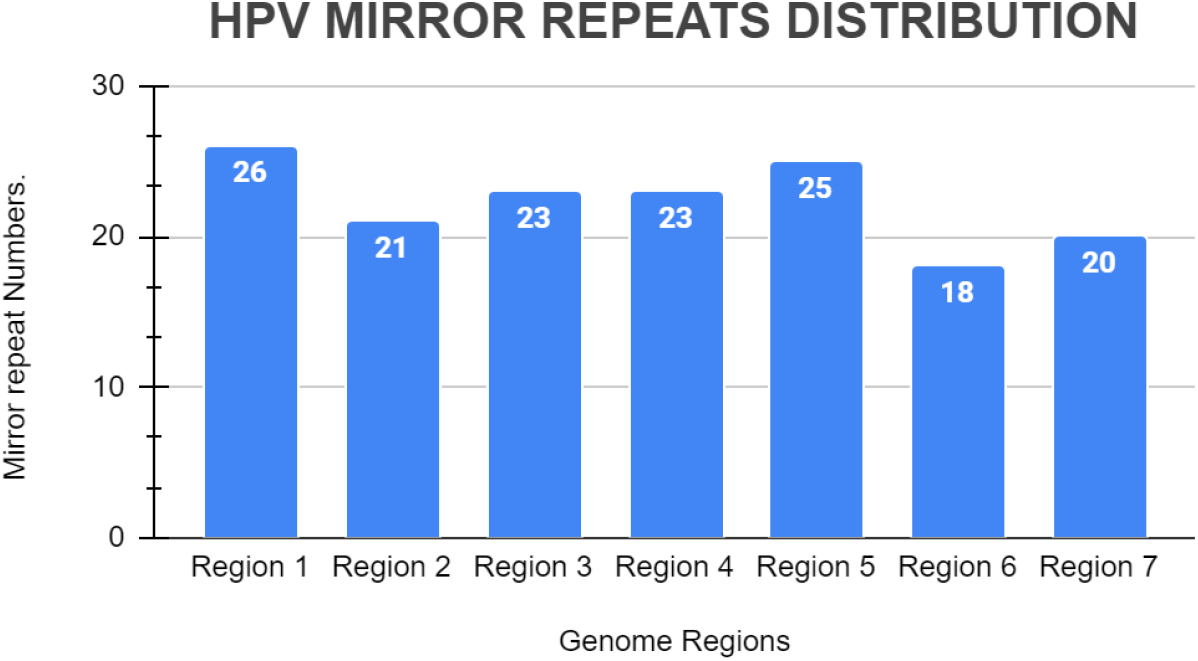
Represent the MR distribution in HPV. The maximum & minimum no of MR sequences were reported in Region 1 (26) & Region 6 (18) respectively.

**Figure 7.**
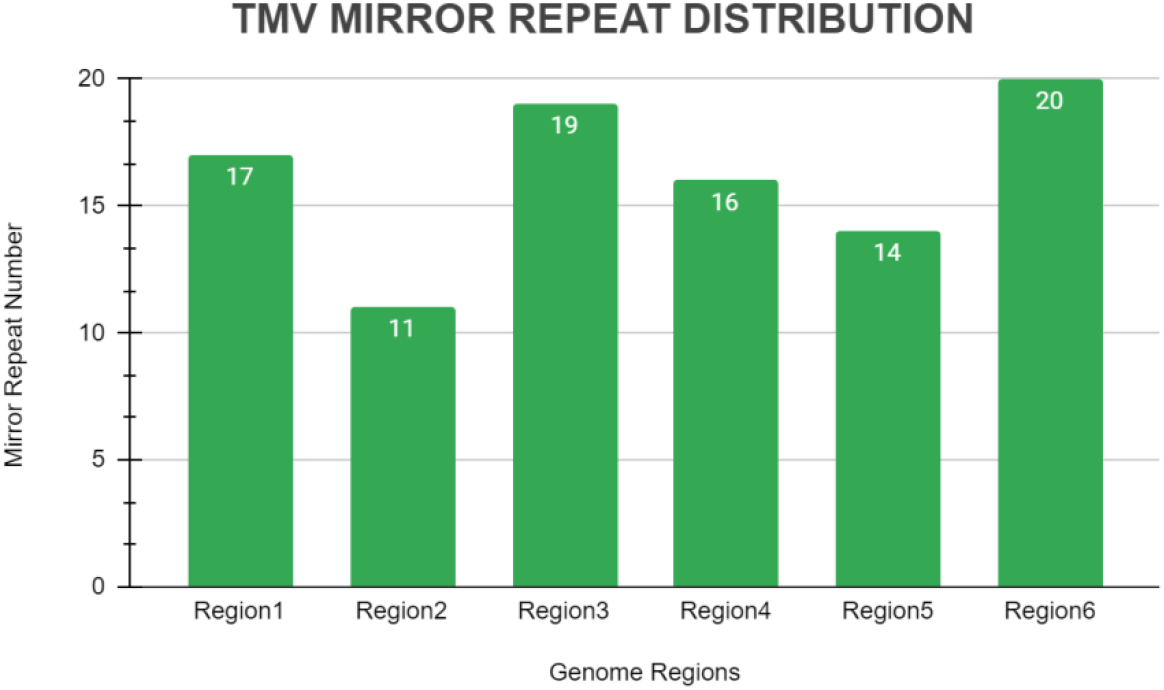
Represent the MR distribution in TMV. The maximum & minimum no of MR sequences were reported in Region 6 (20) & Region 2 (11) respectively.

**Figure 8.**
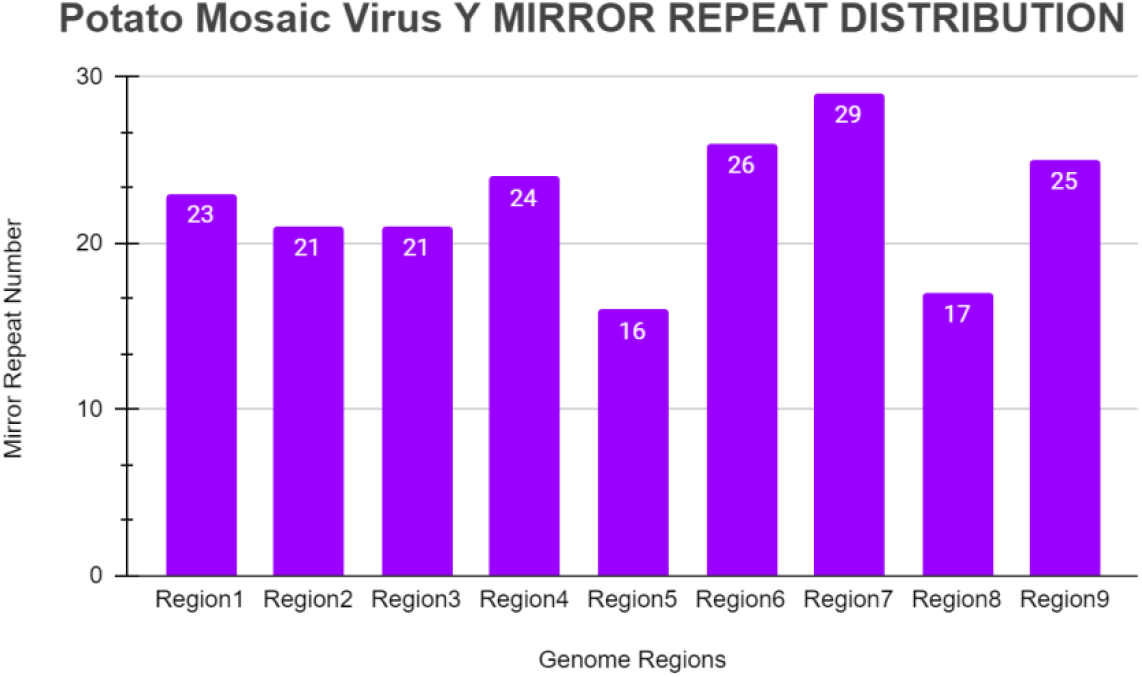
Represent the MR distribution in PMVY. The maximum & minimum no of MR sequences were reported in Region 7 (29) & Region 5 (16) respectively.

By using the said methodology we have identified MR sequences in some selected viral genomes from three different categories (Animal, Plant & Human) given in **Table 1**-

**Table 1.** Shows a list of viruses along with their NCBI reference number

| Serial No. | Name of the Virus | Category | Diseases Association | NCBI Reference No. |
| --- | --- | --- | --- | --- |
| 1 | Rabies Virus | Animal Virus | Rabies | NC_001542.1 |
| 2 | Dengue Virus | Animal Virus | Dengue | NC_001474.2 |
| 3 | HTLV-1 | Human Virus | Cancer | NC_001436.1 |
| 4 | TMV | Plant Virus | Tobacco crop | NC_001367.1 |
| 5 | Potato Virus Y | Plant Virus | Potato crop | NC_001616.1 |
| 6 | HCV | Human Virus | Cancer | NC_004102.1 |
| 7 | HPV | Human Virus | Cancer | NC_027779.1 |

## Result & Discussion

The genomes of the selected viruses were divided into 1000 base pair regions to ensure the maximum identification of Mirror repeats as per the requirements of the bioinformatics approach (FPCB) utilized. The total no of Mirror repeat sequences in the case of selected viruses varies in no and is given below in Table 2. The maximum & minimum no of MR sequences were reported in the case of Dengue virus (229) & TMV (97) respectively. However other viruses (Rabies, HTLV-1, HCV, HPV, PVY) also have a frequent occurrence of MR sequences.

**Table 2.** Represent the total no of Mirror Repeats (MR’s) identified in selected viruses

| Serial No. | Name of the Virus | Category | Total No of MR Sequences |
| --- | --- | --- | --- |
| 1 | Rabies Virus | Animal Virus | 204 |
| 2 | Dengue Virus | Animal Virus | 229 |
| 3 | HTLV-1 | Human Virus | 175 |
| 4 | TMV | Plant Virus | 97 |
| 5 | Potato Virus Y | Plant Virus | 203 |
| 6 | HCV | Human Virus | 178 |
| 7 | HPV | Human Virus | 156 |

The analysis of the results obtained from the approach depicted the occurrence of MR sequences in whole genomic parts of all the selected viruses. The following graphical data represent the region-wise distribution of MR sequences in selected viral genomes.

**Figure 2.**
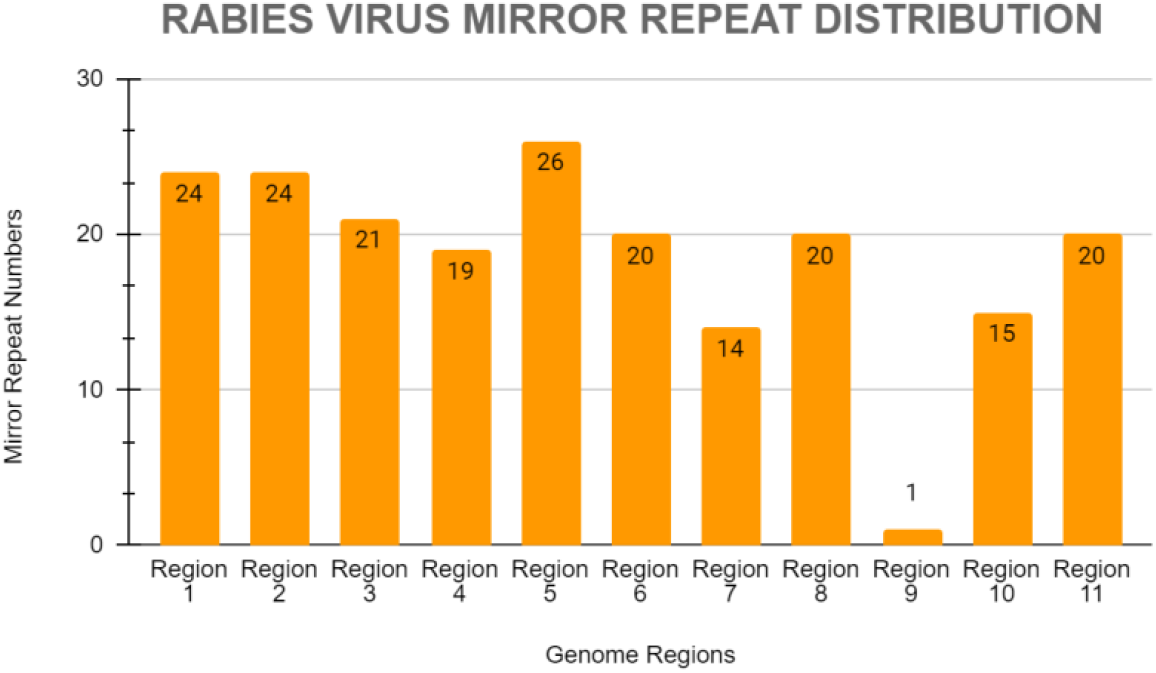
Represent the MR distribution in Rabies virus. The maximum no of MR sequences was reported in Region 5 (26) while the minimum no is in Region 9 (1).

Some of the sequences whose length is found to be more than 21 bps are given below in **Table 3**. [To check the region-wise distribution of MR sequences in all selected viruses including their MR no, Sequences, their length & position in the region, this data provided in the supplementary data file of this manuscript].

**Table 3.**
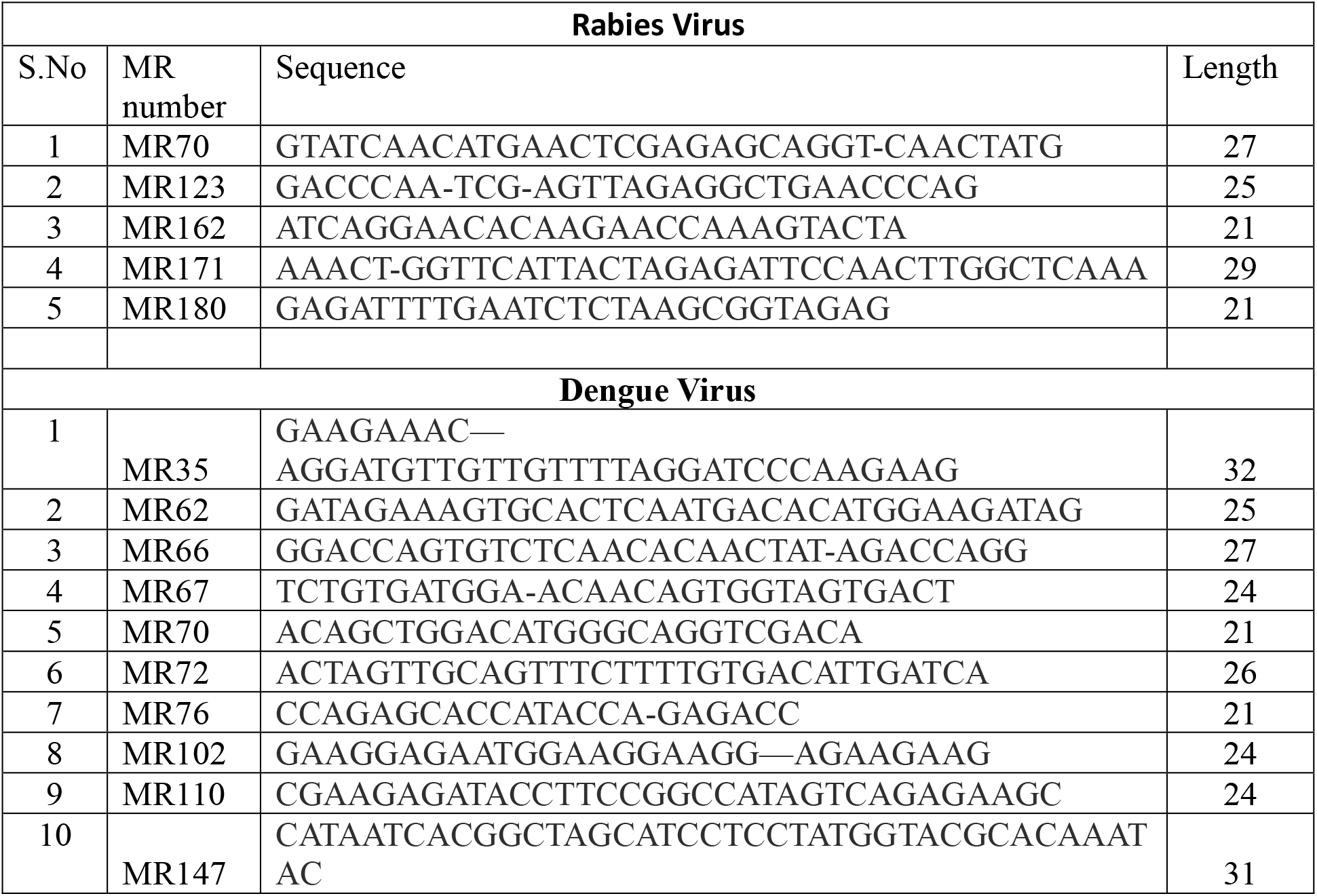
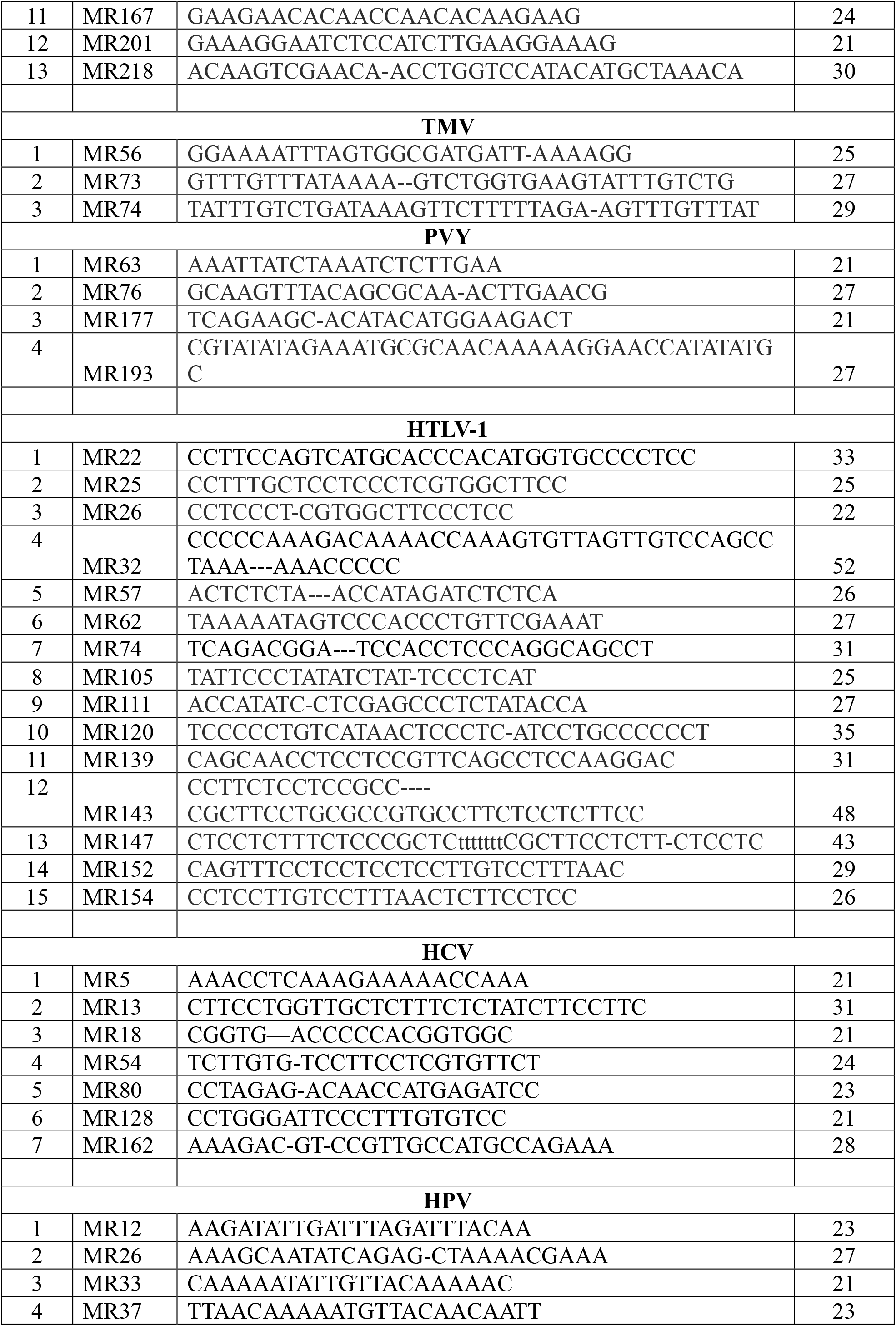
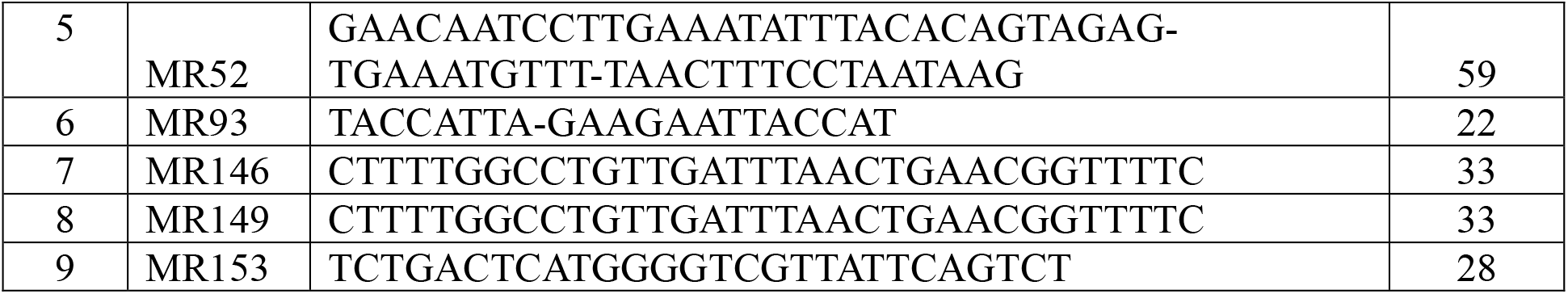
Shows selected sequences & their length in viruses from different categories

### Discussion

The present study is also supported by previous ones in which MR sequences are frequently reported in viruses & various other domains of study by FPCB analysis. The work carried out on HIV identified more than 200 MR sequences with varied lengths in HIV-1 & HIV-2 [23]. The same case is also true in the plant & animal domain [24-27]. The frequent occurrence of mirror repeats correlated with their evolutionary prospective also or may be these sequences show conservative nature in every existing species. *In silico* analysis can be utilized to check their clinical significance as well as their uses in medical science [28-30]. They may be utilized for a novel classification system that is based on mirror repeat profiling. This study will be helpful for computational biology or *in silico*-based studies to find out novel ways of MR identification as well as to study their significant roles & functions in the genes or genomes.

## Conclusion

Based on data obtained from the bioinformatic analysis it is concluded here that the frequent occurrence of mirror repeats in viral genomes indicates their key roles at the genetic level. The observation of short to long sequences within the genomes aids a hint towards their evolutionary as well as clinical significance also.

## Acknowledgment

The authors of this manuscript want to acknowledge the management of Starex University to provide the facility to do the research.

## Author contribution statement

The whole experiments & analysis carried out by Ms. Pooja Yadav, Ms. Priyanka Yadav, Ms. Jyoti Kumari & Ms. Rachna Yadav. Ms. Barkha Sehrawat, Ms. Manisha Yadav, Ms. Amrita Singh, Ms. Shivani Yadav work collectively with the above authors & help in data analysis as well as in the writing of the manuscript. Dr. Dinesh Sharma helps in the drafting & revision of the manuscript. Dr. Sandeep Yadav (Corresponding author) finalized the final version of this manuscript by incorporating the suggestions received from the authors. All authors read & approved the final version of this manuscript.

## Funding Acknowledgment

The present work did not receive any financial support from any funding agency.

## Ethical Statement

This is declared here that in the present work, no method is used unethically.

## References

1. Sanjuán R, Domingo-Calap P. Genetic diversity and evolution of viral populations. Encyclopedia of virology. 2021:53.

2. Cavalli-Sforza LL. The human genome diversity project: past, present and future. Nature Reviews Genetics. 2005 Apr 1;6(4):333 –40.

3. Vellend M, Harmon LJ, Lockwood JL, Mayfield MM, Hughes AR, Wares JP, Sax DF. Effects of exotic species on evolutionary diversification. Trends in Ecology & Evolution. 2007 Sep 1;22(9):481–8.

4. Zamudio KR, Bell RC, Mason NA. Phenotypes in phylogeography: Species’ traits, environmental variation, and vertebrate diversification. Proceedings of the National Academy of Sciences. 2016 Jul 19;113(29):8041 –8.

5. Mager DL, Medstrand P. Retroviral repeat sequences. Nature encyclopedia of the human genome. 2003;5:57–63.

6. Strawbridge EM, Benson G, Gelfand Y, Benham CJ. The distribution of inverted repeat sequences in the Saccharomyces cerevisiae genome. Current genetics. 2010 Aug;56:321–40.

7. Singer MF. Highly repeated sequences in mammalian genomes. International review of cytology. 1982 Jan 1;76:67–112.

8. Qian Z, Adhya S. DNA repeat sequences: diversity and versatility of functions. Current Genetics. 2017 Jun;63:411–6.

9. Todd RT, Wikoff TD, Forche A, Selmecki A. Genome plasticity in Candida albicans is driven by long repeat sequences. Elife. 2019 Jun 7;8:e45954.

10. Duval A, Hamelin R. Mutations at coding repeat sequences in mismatch repair - deficient human cancers: toward a new concept of target genes for instability. Cancer research. 2002 May 1;62(9):2447–54.

11. Liang D, Wilusz JE. Short intronic repeat sequences facilitate circular RNA production. Genes & development. 2014 Oct 15;28(20):2233 –47.

12. Cox R, Mirkin SM. Characteristic enrichment of DNA repeats in different genomes. Proceedings of the National Academy of Sciences. 1997 May 13;94(10):5237 –42.

13. Bigot Y, Brillet B, Auge-Gouillou C. Conservation of palindromic and mirror motifs within inverted terminal repeats of mariner-like elements. Journal of molecular biology. 2005 Aug 5;351(1):108–16.

14. Sagot MF, Viari A. Flexible identification of structural objects in nucleic acid sequences: palindromes, mirror repeats, pseudoknots and triple helices. InCombinatorial Pattern Matching: 8th Annual Symposium, CPM 97 Aarhus, Denmark, June 30–July 2, 1997 Proceedings 8 1997 (pp. 224–246). Springer Berlin Heidelberg.

15. Billingsley KJ, Lättekivi F, Planken A, Reimann E, Kurvits L, Kadastik-Eerme L, Kasterpalu KM, Bubb VJ, Quinn JP, Kõks S, Taba P. Analysis of repetitive element expression in the blood and skin of patients with Parkinson’s disease identifies differential expression of satellite elements. Scientific Reports. 2019 Mar 13;9(1):4369.

16. Mirkin SM, Lyamichev VI, Drushlyak KN, Dobrynin VN, Filippov SA, Frank-Kamenetskii MD. DNA H form requires a homopurine –homopyrimidine mirror repeat. Nature. 1987 Dec 3;330(6147):495–7.

17. Belotserkovskii BP, Veselkov AG, Filippov SA, Dobrynin VN, Mirkin SM, Frank-Kamenetskii MD. Formation of intramolecular triplex in homopurine-homopyrimidine mirror repeats with point substitutions. Nucleic acids research. 1990 Nov 25;18(22):6621–4.

18. Sinden RR. Biological implications of the DNA structures associated with disease-causing triplet repeats. The American Journal of Human Genetics. 1999 Feb 1;64(2):346–53.

19. Bissler JJ. Triplex DNA and human disease. Frontiers in Bioscience-Landmark. 2007 May 1;12(7):4536–46.

20. Kim HM. Genome instability induced by triplex forming mirror repeats in Saccharomyces cerevisiae. Georgia Institute of Technology; 2009

21. Vikash B, Swapni G, Sitaram M, Kulbhushan S. FPCB: a simple and swift strategy for mirror repeat identification. Preprint arXiv: 2013 Dec 13;1312.3869. https://arxiv.org/abs/1312.3869v1

22. . National Center for Biotechnology Information (NCBI)[Internet]. Bethesda (MD): National Library of Medicine (US), National Center for Biotechnology Information; [1988] – [cited 2023 April 2]. Available from: http://www.ncbi.nlm.nih.gov

23. Yadav S, Yadav U, Sharma DC. In-Silico Evaluation of ‘Mirror Repeats’ in HIV Genome.(2021). Int. J. Life Sci. 2022 Sep 7; 11(5): 81–87.

24. Yadav S, Yadav U, Sharma DC. In silico approach for the identification of mirror repeats in selected operon genes of Escherichia coli strain K-12 substrain MG1655. Biomedical and Biotechnology Research Journal (BBRJ). 2022 Jan 1;6(1):93.

25. Yadav U, Yadav S, Sharma DC. Characterization of Flowering Genes of Arabidopsis thaliana for Mirror Repeats. Biointerface Research in Applied Chemistry. 2021;12(3):2852–61.

26. Yadav U, Yadav S, Sharma DC. In Silico Analysis of Structural Photosynthetic Genes of Arabidopsis thaliana for Unique Mirror Repeats. Indian Journal of Science and Technology. 2022 Feb 4;15(3):127–35.

27. Yadav D, Dhankhar M, Saini K, Bhardwaj V. A novel approach for identification of mirror repeats within the Engrailed Homeobox-1 gene of Xenopus tropicalis. Biomedical and Biotechnology Research Journal (BBRJ). 2022 Oct 1;6(4):532.

28. Mondal SI, Ferdous S, Jewel NA, Akter A, Mahmud Z, Islam MM, Afrin T, Karim N. Identification of potential drug targets by subtractive genome analysis of Escherichia coli O157: H7: an in silico approach. Advances and applications in bioinformatics and chemistry. 2015 Dec 8:49–63.

29. Yazdani Z, Rafiei A, Yazdani M, Valadan R. Design an efficient multi-epitope peptide vaccine candidate against SARS-CoV-2: an in silico analysis. Infection and drug resistance. 2020 Aug 25:3007–22.

30. Farhadi T. Effectiveness assessment of protein drugs and vaccines through in Silico analysis. Biomedical and Biotechnology Research Journal (BBRJ). 2018 Apr 1;2(2):106.

