## Supplementary Data File for "Identification of Mirror Repeats in Viral Genomes using FPCB Analysis"

HCV VIRUS

| | | | | |
|---|---|---|---|---|
| REGION 1 | | | | |
| MIRROR REPEATS | SEQUENCE | LENGTH | POSITION IN REGION | Total |
| MR1 | GTTAGTATGAGTG | 13 | 90 | |
| MR2 | GCCCCCG | 7 | 235 | |
| MR3 | AGCACGA | 7 | 345 | |
| MR4 | AATCCTAA | 8 | 351 | |
| MR5 | AAACCTCAAAGAAAAACCAAA | 21 | 357 | |
| MR6 | TGGCGGT | 7 | 419 | |
| MR7 | GCGCGCG | 7 | 479 | 15 |
| MR8 | CCTATCC | 7 | 531 | |
| MR9 | GCGGGTGGGCG | 11 | 613 | |
| MR10 | GGGCGGG | 7 | 619 | |
| MR11 | GGCTCTCGG | 9 | 645 | |
| MR12 | CATGGGGTAC | 10 | 740 | |
| MR13 | CTTCCTGGTTGCTCTTTCTCTATCTTCCTTC | 31 | 846 | |
| MR14 | TCTTCCTTCT | 10 | 868 | |
| MR15 | TCGGGGCT | 8 | 933 | |
| REGION 2 | | | | |
| MR16 | CTTGCGTTC | 9 | 24 | |
| MR17 | GTGGCGGTG | 9 | 59 | |
| MR18 | CGGTG--ACCCCCACGGTGGC | 21 | 63 | |
| MR19 | CAGGGAC | 7 | 85 | |
| MR20 | GCAGCTTCGACG | 12 | 109 | |
| MR21 | TCTGTCT | 7 | 188 | |
| MR22 | TCTTTCT | 7 | 192 | |
| MR23 | TTGGTCAACTGTTT | 14 | 201 | |
| MR24 | TCTATCT | 7 | 260 | 20 |
| MR25 | CTATCTATC | 9 | 261 | |
| MR26 | CGTTGGTGGTAGC | 13 | 333 | |
| MR27 | GAAGGTCCTGGTAG | 14 | 448 | |
| MR28 | GCCGGCCG | 8 | 515 | |
| MR29 | GCACCACG | 8 | 522 | |
| MR30 | CTGGGGTC | 8 | 745 | |
| MR31 | GTGTGTG | 7 | 845 | |
| MR32 | TGGCCCGGT | 9 | 850 | |
| MR33 | GTGGTGGTG | 9 | 881 | |
| MR34 | GGTGGTGG | 8 | 883 | |
| MR35 | GCCACCG | 7 | 970 | |
| REGION 3 | | | | |
| MR36 | AACTCAA | 7 | 7 | |
| MR37 | GGGTGGG | 7 | 59 | |
| MR38 | CAACAAC | 7 | 66 | |
| MR39 | GGTCCCTGG | 9 | 139 | |
| MR40 | GGGAGGG | 7 | 240 | |
| MR41 | GCGGGGCG | 8 | 282 | |
| MR42 | GACAGGGACAG | 11 | 307 | |
| MR43 | GGACAGG | 7 | 312 | |
| MR44 | TCCACCT | 7 | 410 | |
| MR45 | CCACCTCCACC | 11 | 411 | |
| MR46 | GACGTGCAG | 9 | 433 | |
| MR47 | TCCTCCT | 7 | 509 | |
| MR48 | GCAGACG | 7 | 520 | |
| MR49 | CGCGCGC | 7 | 525 | |
| MR50 | GCGCGCG | 7 | 526 | 27 |
| MR51 | CGTCTGC | 7 | 531 | |
| MR52 | TACTCAT | 7 | 557 | |
| MR53 | GCGGAGGCG | 9 | 571 | |
| MR54 | TCTTGTG-TCCTTCCTCGTGTTCT | 24 | 633 | |
| MR55 | GTGGGTG | 7 | 684 | |
| MR56 | CCTC-TCCTCCTGCTCC | 17 | 724 | |
| MR57 | GCATACG | 7 | 760 | |
| MR58 | TGTTCTTGT | 9 | 804 | |
| MR59 | CTACATC | 7 | 852 | |

| | | | | | | |
|---|---|---|---|---|---|---|
| MR60 | | TGCACGT | | 7 | 911 | |
| MR61 | | TACTCAT | | 7 | 965 | |
| MR62 | | TGTGTGT | | 7 | 971 | |
| REGION 4 | | | | | | |
| MR63 | | TGCGCGT | | 7 | 79 | |
| MR64 | | ACGTGCA | | 7 | 139 | |
| MR65 | | TATGTGTAT | | 9 | 183 | |
| MR66 | | GCGTGCG | | 7 | 315 | |
| MR67 | | TGCCCGTCTCTGCCCGT | | 17 | 340 | |
| MR68 | | CAGCCGAC | | 8 | 382 | |
| MR69 | | TGGAGGT | | 7 | 408 | |
| MR70 | | AGCAGACGA | | 9 | 442 | |
| MR71 | | CAAAAAC | | 7 | 494 | 19 |
| MR72 | | GTCAACTG | | 8 | 527 | |
| MR73 | | CCAAACC | | 7 | 539 | |
| MR74 | | ACGTGCA | | 7 | 555 | |
| MR75 | | ATGGGGTA | | 8 | 565 | |
| MR76 | | GGCTCCTCGG | | 10 | 717 | |
| MR77 | | CCTGCTTTCGCCC | | 13 | 794 | |
| MR78 | | CCCGGCCC | | 8 | 805 | |
| MR79 | | GGCTCCTCGG | | 10 | 828 | |
| MR80 | | CCTAGAG-ACAACCATGAGATCC | | 23 | 941 | |
| MR81 | | CCTCTCC | | 7 | 982 | |
| REGION 5 | | | | | | |
| MR82 | | TCCACCT | | 7 | 221 | |
| MR83 | | GTTCCTTG | | 8 | 235 | |
| MR84 | | TAATAAT | | 7 | 276 | |
| MR85 | | CAGAGAC | | 7 | 348 | |
| MR86 | | GCGGGGGCG | | 9 | 356 | |
| MR87 | | GTCACTG | | 7 | 404 | |
| MR88 | | CTGTGTC | | 7 | 408 | |
| MR89 | | CCACCACC | | 8 | 444 | |
| MR90 | | TCCCCCT | | 7 | 480 | 23 |
| MR91 | | AAGGGGGGAA | | 10 | 497 | |
| MR92 | | AAGAAGAA | | 8 | 530 | |
| MR93 | | GAAGAAG | | 7 | 532 | |
| MR94 | | GCTGGTCG | | 8 | 559 | |
| MR95 | | TGTCTGT | | 7 | 609 | |
| MR96 | | CGACCAGC | | 8 | 621 | |
| MR97 | | AGCGGCGA | | 8 | 626 | |
| MR98 | | TTACCATT | | 8 | 750 | |
| MR99 | | AGGGGGA | | 7 | 818 | |
| MR100 | | AGGCATCTACAGA | | 13 | 829 | |
| MR101 | | GTGAGTG | | 7 | 894 | |
| MR102 | | CGCCCGC | | 7 | 933 | |
| MR103 | | GTACATG | | 7 | 964 | |
| MR104 | | CCGTGTGCC | | 9 | 987 | |
| REGION 6 | | | | | | |
| MR105 | | GGGAGGG | | 7 | 14 | |
| MR106 | | CTCACTC | | 7 | 34 | |
| MR107 | | GCCACCG | | 7 | 112 | |
| MR108 | | CGTGTGC | | 7 | 117 | |
| MR109 | | CCCCTCCCC | | 9 | 137 | |
| MR110 | | TCCGCCT | | 7 | 176 | |
| MR111 | | CCCACCC | | 7 | 187 | |
| MR112 | | CTGGAGGTC | | 9 | 298 | |
| MR113 | | CACGAGCAC | | 9 | 309 | |
| MR114 | | GGGTGCTCGTTGG | | 13 | 320 | 18 |
| MR115 | | CCAGACC | | 7 | 591 | |
| MR116 | | GGTCTTCTGG | | 10 | 615 | |
| MR117 | | CTGCCGTC | | 8 | 725 | |
| MR118 | | CCAAACC | | 7 | 753 | |
| MR119 | | ggTGGGTGG | | 9 | 782 | |
| MR120 | | GCCCCCG | | 7 | 805 | |
| MR121 | | GGTATGG | | 7 | 902 | |
| MR122 | | GGCGCGGGCGTGG | | 13 | 907 | |

| | | | | |
|---|---|---|---|---|
| REGION 7 | | | | |
| MR123 | GGTGTGG | 7 | 24 | |
| MR124 | GGCCCGG | 7 | 60 | |
| MR125 | CACGCAC | 7 | 134 | |
| MR126 | CGCCCGC | 7 | 164 | |
| MR127 | CGTCACTGC | 9 | 170 | |
| MR128 | CCTGGGATTCCCTTTGTGTCC | 21 | 351 | 17 |
| MR129 | GGTCTGG | 7 | 392 | |
| MR130 | CAAAAAC | 7 | 458 | |
| MR131 | TCGCGCT | 7 | 580 | |
| MR132 | GGGTGGG | 7 | 625 | |
| MR133 | TGCCCGT | 7 | 675 | |
| MR134 | CCGTGCC | 7 | 678 | |
| MR135 | GGAGGAGG | 8 | 767 | |
| MR136 | AGCCCGA | 7 | 829 | |
| MR137 | CCCTCCC | 7 | 873 | |
| MR138 | CCAGCTCCTCGGCC | 14 | 937 | |
| MR139 | ACCGCCA | 7 | 987 | |
| REGION 8 | | | | |
| MR140 | AGTGGTGA | 8 | 88 | |
| MR141 | AGGAGGA | 7 | 126 | |
| MR142 | CCCGGGCCC | 9 | 186 | |
| MR143 | TGCCCGT | 7 | 195 | |
| MR144 | GGCGCGG | 7 | 205 | |
| MR145 | GAAAAAG | 7 | 244 | |
| MR146 | CACCTCCAC | 9 | 291 | |
| MR147 | GCCTCCG | 7 | 313 | |
| MR148 | CTCCGCCTC | 9 | 315 | |
| MR149 | GAAAAAG | 7 | 325 | |
| MR150 | GGCATTACGG | 10 | 416 | |
| MR151 | CCCGCCC | 7 | 452 | 23 |
| MR152 | GAGGGGGAG | 9 | 515 | |
| MR153 | CAGCGAC | 7 | 541 | |
| MR154 | GTCGTGTGCTG | 11 | 590 | |
| MR155 | CTTATTC | 7 | 609 | |
| MR156 | GGACAGG | 7 | 618 | |
| MR157 | CAAAAAC | 7 | 656 | |
| MR158 | TCGTTGCT | 8 | 686 | |
| MR159 | CACTTCAC | 8 | 721 | |
| MR160 | CCATTACC | 8 | 787 | |
| MR161 | AAGTGAA | 7 | 831 | |
| MR162 | AAAGAC-GT-CCGTTGCCATGCCAGAAA | 28 | 917 | |
| REGION 9 | | | | |
| MR163 | ACTACCATCA | 10 | 7 | |
| MR164 | TGGGCGTGCGCGT | 13 | 95 | |
| MR165 | GTGCGCGTG | 9 | 100 | |
| MR166 | GCGTGTGCG | 9 | 104 | |
| MR167 | AGTCACTGA | 9 | 282 | |
| MR168 | ACGGAGGAGGCA | 12 | 304 | |
| MR169 | AAGGGGGGAA | 10 | 408 | |
| MR170 | GTGAAAGTG | 9 | 572 | 16 |
| MR171 | CAGGAGGAC | 9 | 589 | |
| MR172 | ATTTCTTTA | 9 | 884 | |
| MR173 | TCCCGCCCT | 9 | 1082 | |
| MR174 | CTCAAACTC | 9 | 1201 | |
| MR175 | AAACTCACTCCAA | 13 | 1204 | |
| MR176 | CCCGGCCC | 8 | 1301 | |
| MR177 | CCTCCTCC | 8 | 1359 | |
| MR178 | CTCCGGCCTC | 10 | 1393 | |
| | HPV | | | |
| REGION 1 | | | | |
| MIRROR REPEATS | SEQUENCE | LENGTH | POSITION IN REGION | TOTAL |
| MR1 | CAAGAAC | 7 | 17 | |
| MR2 | CAGTTTGAC | 9 | 43 | |
| MR3 | AACTATCAA | 9 | 103 | |
| MR4 | TTCTCTT | 7 | 126 | |
| MR5 | TTAGAGATT | 9 | 152 | |

| | | | | | | |
|---|---|---|---|---|---|---|
| MR6 | | GTATTATG | | 8 | 165 | |
| MR7 | | GTTTGAGTTTG | | 11 | 207 | |
| MR8 | | GAGTTTGAG | | 9 | 211 | |
| MR9 | | GAAAAAG | | 7 | 268 | |
| MR10 | | TGAAAAGT | | 8 | 330 | 26 |
| MR11 | | TGTTTGT | | 7 | 345 | |
| MR12 | | AAGATATTGATTTAGATTTACAA | | 23 | 442 | |
| MR13 | | ATTTAGATTTA | | 11 | 451 | |
| MR14 | | GAGGAGGAG | | 9 | 528 | |
| MR15 | | TTATATT | | 7 | 543 | |
| MR16 | | TTGGGTT | | 7 | 548 | |
| MR17 | | GTGTGTG | | 7 | 591 | |
| MR18 | | TGTGTGT | | 7 | 592 | |
| MR19 | | GTTTTTG | | 7 | 597 | |
| MR20 | | ATTTTTA | | 7 | 632 | |
| MR21 | | TTTCCTTT | | 8 | 652 | |
| MR22 | | GAAAAAG | | 7 | 733 | |
| MR23 | | TGATAGT | | 7 | 741 | |
| MR24 | | ATTGTTA | | 7 | 754 | |
| MR25 | | TTAAATT | | 7 | 784 | |
| MR26 | | AAAGCAATATCAGAG-CTAAAACGAAA | | 27 | 919 | |
| REGION 2 | | | | | | |
| MR27 | | TTAAAATT | | 8 | 1 | |
| MR28 | | AAGGAAAAAGTAA | | 13 | 16 | |
| MR29 | | GATGTAG | | 7 | 148 | |
| MR30 | | AAATTTAAA | | 9 | 180 | |
| MR31 | | ACAAAACA | | 8 | 238 | |
| MR32 | | GAAGAAG | | 7 | 285 | |
| MR33 | | CAAAAATATTGTTACAAAAAC | | 21 | 304 | |
| MR34 | | TACAAAAACAT | | 11 | 316 | 21 |
| MR35 | | TTATATT | | 7 | 366 | |
| MR36 | | TTGAGTT | | 7 | 376 | |
| MR37 | | TTAACAAAAATGTTACAACAATT | | 23 | 408 | |
| MR38 | | AATTATTAA | | 9 | 427 | |
| MR39 | | TTATATT | | 7 | 495 | |
| MR40 | | ATATTTTTATA | | 11 | 497 | |
| MR41 | | AATATAA | | 7 | 522 | |
| MR42 | | ATTGGTTA | | 8 | 553 | |
| MR43 | | AGTCTTTCTGA | | 11 | 609 | |
| MR44 | | AAACGCAAA | | 9 | 698 | |
| MR45 | | CGCAAACGC | | 9 | 701 | |
| MR46 | | CATCCTAC | | 8 | 835 | |
| MR47 | | AAATGTAAA | | 9 | 890 | |
| REGION 3 | | | | | | |
| MR48 | | TTAAATT | | 7 | 14 | |
| MR49 | | AAGTCATTTCTTTCATGAA | | 19 | 33 | |
| MR50 | | TTCACTT | | 7 | 144 | |
| MR51 | | TATCTAT | | 7 | 177 | |
| MR52 | | GAACAATCCTTGAAATATTTACACAGTAGAG-TGAAATGTTT-TAACTTTCCTAATAAG | | 59 | 260 | |
| MR53 | | CAACAAC | | 7 | 321 | |
| MR54 | | GTTATTG | | 7 | 526 | |
| MR55 | | AGAAAAGA | | 8 | 543 | |
| MR56 | | ATTATATTA | | 9 | 560 | |
| MR57 | | AGAAAAGA | | 8 | 573 | |
| MR58 | | CCTGTCC | | 7 | 606 | 23 |
| MR59 | | CCCAACCC | | 8 | 611 | |
| MR60 | | AAAGAAA | | 7 | 699 | |
| MR61 | | ATAAATA | | 7 | 738 | |
| MR62 | | AGTATGGTATGA | | 12 | 788 | |
| MR63 | | ATACCCATA | | 9 | 818 | |
| MR64 | | CATATAC | | 7 | 823 | |
| MR65 | | GGAAAAGG | | 8 | 835 | |
| MR66 | | CAATTAAC | | 8 | 873 | |
| MR67 | | GAAGGAAAGGTAG | | 13 | 882 | |
| MR68 | | AAAGGTAGATGAAAA | | 15 | 887 | |
| MR69 | | TTTATTT | | 7 | 937 | |
| MR70 | | TATTTTAT | | 8 | 939 | |

| | | | | | | |
|---|---|---|---|---|---|---|
| REGION 4 | | | | | | |
| MR71 | | ATAGAGATA | | 9 | 3 | |
| MR72 | | TCTCCTCT | | 8 | 21 | |
| MR73 | | CCTTCACCCAACCCTCTACC | | 20 | 66 | |
| MR74 | | GGAGGAGG | | 8 | 209 | |
| MR75 | | AGGAGGA | | 7 | 211 | |
| MR76 | | TTCCCTT | | 7 | 234 | |
| MR77 | | ACAGGACA | | 8 | 311 | |
| MR78 | | TTCAGCGACTT | | 11 | 351 | |
| MR79 | | AATTTTAA | | 8 | 391 | |
| MR80 | | AATAGAAATAAACATAA | | 17 | 431 | |
| MR81 | | GGGAGGG | | 7 | 492 | 23 |
| MR82 | | ATACATA | | 7 | 501 | |
| MR83 | | ACATACA | | 7 | 503 | |
| MR84 | | AAGAGAA | | 7 | 544 | |
| MR85 | | TGTAAAATGT | | 10 | 615 | |
| MR86 | | AAATATCTATCAA | | 13 | 659 | |
| MR87 | | TGATGTAGT | | 9 | 701 | |
| MR88 | | AAATAAA | | 7 | 710 | |
| MR89 | | AGCAAACGA | | 9 | 721 | |
| MR90 | | ATTATTA | | 7 | 743 | |
| MR91 | | TAAAAAT | | 7 | 748 | |
| MR92 | | TTTATTT | | 7 | 769 | |
| MR93 | | TACCATTA-GAAGAATTACCAT | | 22 | 877 | |
| REGION 5 | | | | | | |
| MR94 | | ATGGGTA | | 7 | 69 | |
| MR95 | | GTAGATG | | 7 | 92 | |
| MR96 | | ACTGGTCA | | 8 | 122 | |
| MR97 | | CATCCTAC | | 8 | 128 | |
| MR98 | | GATGTAG | | 7 | 155 | |
| MR99 | | TATCTAT | | 7 | 237 | |
| MR100 | | TGACAGT | | 7 | 247 | |
| MR101 | | CAACAAC | | 7 | 264 | |
| MR102 | | AACTATCAA | | 9 | 274 | |
| MR103 | | AATGTTTTTGTAA | | 13 | 281 | |
| MR104 | | GAATTAAG | | 8 | 332 | |
| MR105 | | ATAAATA | | 7 | 344 | |
| MR106 | | ATTTTTA | | 7 | 407 | 25 |
| MR107 | | TTTATTT | | 7 | 511 | |
| MR108 | | AGTATGA | | 7 | 519 | |
| MR109 | | GCCTTTTCCG | | 10 | 533 | |
| MR110 | | ACTTTCA | | 7 | 602 | |
| MR111 | | GATTTAG | | 7 | 666 | |
| MR112 | | ATTTTTA | | 7 | 735 | |
| MR113 | | TCAGGACT | | 8 | 796 | |
| MR114 | | GTAGATG | | 7 | 824 | |
| MR115 | | TTGAAGAGAATTT | | 13 | 864 | |
| MR116 | | AAGTTTTTGAA | | 11 | 909 | |
| MR117 | | AAACTGAGGAGGA-TCAAA | | 19 | 945 | |
| MR118 | | GACTCAG | | 7 | 986 | |
| REGION 6 | | | | | | |
| MR119 | | TCCTCCT | | 7 | 54 | |
| MR120 | | AGTAGATGA | | 9 | 66 | |
| MR121 | | ATTCAGACTTA | | 11 | 164 | |
| MR122 | | ACCACCA | | 7 | 230 | |
| MR123 | | GTTGTTG | | 7 | 276 | |
| MR124 | | ATTATTA | | 7 | 317 | |
| MR125 | | GTCTCTG | | 7 | 390 | |
| MR126 | | AATATATAA | | 9 | 464 | |
| MR127 | | AGAAAGGGAACGA | | 13 | 476 | 18 |
| MR128 | | GACACAG | | 7 | 579 | |
| MR129 | | AATCCTAA | | 8 | 588 | |
| MR130 | | ACAAACA | | 7 | 611 | |
| MR131 | | TTGTCCACCTATT | | 13 | 746 | |
| MR132 | | GACACAG | | 7 | 798 | |
| MR133 | | AACAGGACAA | | 10 | 835 | |
| MR134 | | GATATAG | | 7 | 861 | |
| MR135 | | TAAAAAT | | 7 | 898 | |

| | | | | | |
|---|---|---|---|---|---|
| MR136 | | AGAAGTATATGGAGA | 15 | 911 | |
| REGION 7 | | | | | |
| MR137 | | AGTCCTGA | 8 | 44 | |
| MR138 | | ATTGGTTA | 8 | 159 | |
| MR139 | | AATAATAA | 8 | 203 | |
| MR140 | | ACAGTAGTTGATAACA | 16 | 221 | |
| MR141 | | AAACACAAA | 9 | 241 | |
| MR142 | | ACTATATCA | 9 | 254 | |
| MR143 | | TATGATGTAGAAT | 13 | 353 | |
| MR144 | | ACCGCCGCCA | 10 | 466 | |
| MR145 | | TATAGATAT | 9 | 497 | |
| MR146 | | CTTTTGGCCTGTTGATTTAACTGAACGGTTTTC | 33 | 580 | |
| MR147 | | CTGAATTAAGTC | 12 | 615 | 20 |
| MR148 | | CCTGGGTCC | 9 | 634 | |
| MR149 | | CTTTTGGCCTGTTGATTTAACTGAACGGTTTTC | 33 | 679 | |
| MR150 | | CTGAATTAAGTC | 12 | 714 | |
| MR151 | | GCAAAACG | 8 | 836 | |
| MR152 | | TAAAATAATAATAT | 14 | 852 | |
| MR153 | | TCTGACTCATGGGGTCGTTATTCAGTCT | 28 | 931 | |
| MR154 | | GTATATATG | 9 | 967 | |
| MR155 | | AAGATTTTTAAAA | 13 | 1108 | |
| MR156 | | TACAACAT | 8 | 1123 | |
| | | DENGUE VIRUS | | | |
| REGION 1 | | | | | |
| MIRROR REPEATS | | SEQUENCE | LENGTH | POSITION IN REGION | TOTAL |
| MR1 | | AGACAGA | 7 | 27 | |
| MR2 | | GAGGGAG | 7 | 40 | |
| MR3 | | CGGAAAAAGGC | 11 | 109 | |
| MR4 | | GCAGGGACG | 9 | 210 | |
| MR5 | | CCTAACAATCC | 11 | 264 | |
| MR6 | | AATTAAAAAATCAA | 12 | 309 | |
| MR7 | | AGACGCAGA | 9 | 388 | |
| MR8 | | AAGAGAA | 7 | 488 | |
| MR9 | | AAAGGGAAA | 9 | 493 | |
| MR10 | | GTCTTCTG | 8 | 503 | |
| MR11 | | CATGTGTAC | 9 | 534 | 17 |
| MR12 | | CAGAAGAC | 8 | 620 | |
| MR13 | | GTACCACCATG | 11 | 677 | |
| MR14 | | AAGAGAA | 7 | 699 | |
| MR15 | | AGAAAAAAGA | 10 | 702 | |
| MR16 | | GACTTTGTGGAAGGGGTTTCAG | 20 | 964 | |
| MR17 | | AGGAGGA | 7 | 984 | |
| REGION 2 | | | | | |
| MR18 | | CAAAAAAC | 8 | 40 | |
| MR19 | | AAACAAA | 7 | 44 | |
| MR20 | | CAACACAAC | 9 | 134 | |
| MR21 | | AACACAACAACAGAA | 13 | 135 | |
| MR22 | | CCAACACAAGGGGAACCCAGCC | 18 | 159 | |
| MR23 | | CGTCTGC | 7 | 206 | |
| MR24 | | AGACAGA | 7 | 227 | |
| MR25 | | GGAAAGG | 7 | 261 | |
| MR26 | | CTCACTC | 7 | 364 | |
| MR27 | | AAATGACACAGGAAA | 13 | 392 | |
| MR28 | | ACACCACA | 8 | 429 | 22 |
| MR29 | | CACTGTCAC | 9 | 473 | |
| MR30 | | CAAGAAC | 7 | 496 | |
| MR31 | | AAATAAA | 7 | 542 | |
| MR32 | | ACACACA | 7 | 610 | |
| MR33 | | AGAAAGA | 7 | 634 | |
| MR34 | | ACTTTCA | 7 | 651 | |
| MR35 | | GAAGAAAC--AGGATGTTGTTGTTTTAGGATCCCAAGAAG | 32 | 671 | |
| MR36 | | ACTCTTCACAGGACA--TCTCA | 18 | 767 | |
| MR37 | | ATGTCATACTCTA | 11 | 825 | |

| | | | | | | |
|---|---|---|---|---|---|---|
| MR38 | | AACACAA | | 7 | 878 | |
| MR39 | | GAAAAAAG | | 8 | 963 | |
| REGION 3 | | | | | | |
| MR40 | | GACAGAAAAAGATAG | | 13 | 10 | |
| MR41 | | ACCTCCA | | 7 | 46 | |
| MR42 | | GGAGACAGCTACATCATCATAGG | | 13 | 56 | |
| MR43 | | AAGAGAA | | 7 | 164 | |
| MR44 | | GACACAG | | 7 | 185 | |
| MR45 | | CATCTATAGGAAAGGCTCTCCAC | | 17 | 225 | |
| MR46 | | GGATAGG | | 7 | 330 | |
| MR47 | | CACTGTCTGTGAC | | 11 | 357 | |
| MR48 | | GTTGCGTTG | | 9 | 429 | |
| MR49 | | AAACAAA | | 7 | 448 | |
| MR50 | | ACAGACA | | 7 | 485 | |
| MR51 | | ACGTGCA | | 7 | 492 | 21 |
| MR52 | | TCAAAACT | | 8 | 539 | |
| MR53 | | AAACAAA | | 7 | 626 | |
| MR54 | | CCAGACC | | 7 | 811 | |
| MR55 | | AACACAA | | 7 | 854 | |
| MR56 | | GAAGTTGAAG | | 10 | 881 | |
| MR57 | | ACCACCA | | 7 | 911 | |
| MR58 | | AAAGAAA | | 7 | 935 | |
| MR59 | | ACTCAAAACTCA | | 12 | 960 | |
| MR60 | | AGACAACAGA | | 10 | 988 | |
| REGION 4 | | | | | | |
| MR61 | | ATGGGTTATTGGATA | | 13 | 13 | |
| MR62 | | GATAGAAAGTGCACTCAATGACACATGGAAGATAG | | 25 | 24 | |
| MR63 | | AGATAGA | | 7 | 53 | |
| MR64 | | ATTGAAGTTA | | 10 | 73 | |
| MR65 | | CACACAC | | 7 | 104 | |
| MR66 | | GGACCAGTGTCTCAACACAACTAT-AGACCAGG | | 27 | 166 | |
| MR67 | | TCTGTGATGGA-ACAACAGTGGTAGTGACT | | 24 | 257 | |
| MR68 | | AGGAGAAAGAAGA | | 11 | 437 | |
| MR69 | | AAGAGAA | | 7 | 446 | |
| MR70 | | ACAGCTGGACATGGGCAGGTCGACA | | 21 | 472 | |
| MR71 | | ACTTTTCA | | 8 | 497 | |
| MR72 | | ACTAGTTGCAGTTTCTTTTGTGACATTGATCA | | 26 | 579 | 20 |
| MR73 | | GGTATGG | | 7 | 682 | |
| MR74 | | ACCTCCA | | 7 | 766 | |
| MR75 | | CCTCTCC | | 7 | 807 | |
| MR76 | | CCAGAGCACCATACCA-GAGACC | | 21 | 813 | |
| MR77 | | TACCAGAGACCAT | | 13 | 824 | |
| MR78 | | TTGAGTT | | 7 | 839 | |
| MR79 | | CAGTGAC | | 7 | 914 | |
| MR80 | | ACTATCA | | 7 | 919 | |
| REGION 5 | | | | | | |
| MR81 | | CAAAAAAC | | 8 | 36 | |
| MR82 | | CAAGAAC | | 7 | 112 | |
| MR83 | | AAAAATAT | | 8 | 196 | |
| MR84 | | GAGAGAG | | 7 | 291 | |
| MR85 | | CAATAAC | | 7 | 355 | |
| MR86 | | ATAACAATA | | 9 | 357 | |
| MR87 | | GATGGTAG | | 8 | 372 | 25 |
| MR88 | | TAAAAAAT | | 8 | 388 | |
| MR89 | | ACAAACA | | 7 | 407 | |
| MR90 | | TACTCAT | | 7 | 421 | |
| MR91 | | TCAGGACT | | 8 | 450 | |
| MR92 | | CTTTTTC | | 7 | 456 | |
| MR93 | | CATGGTAC | | 8 | 490 | |
| MR94 | | GAAGTGAAG | | 9 | 504 | |
| MR95 | | CACCCCCAC | | 9 | 547 | |
| MR96 | | CCCACCC | | 7 | 551 | |

| | | | | | |
|---|---|---|---|---|---|
| MR97 | | GGAAAGG | 7 | 561 | |
| MR98 | | GAATTAAG | 8 | 592 | |
| MR99 | | AAAGGAAA | 8 | 702 | |
| MR100 | | TCAAGAAAGACCT | 11 | 736 | |
| MR101 | | GGAGGAGG | 8 | 759 | |
| MR102 | | GAAGGAGAATGGAAGGAAGG--AGAAGAAG | 24 | 777 | |
| MR103 | | GTCCAAACGAAACCTG | 14 | 846 | |
| MR104 | | AAAAGGAAAA | 10 | 947 | |
| MR105 | | TTATGGTAATGGTGTT | 14 | 968 | |
| REGION 6 | | | | | |
| MR106 | | ATAT-GTGAGTGCTATA | 15 | 1 | |
| MR107 | | GAAAAAAG | 8 | 26 | |
| MR108 | | AAGAGAA | 7 | 74 | |
| MR109 | | ACCTCCA | 7 | 96 | |
| MR110 | | CGAAGAGATACCTTCCGGCCATAGTCAGAGAAGC | 24 | 120 | |
| MR111 | | TACCAGACCCCAG-CCAT | 16 | 245 | |
| MR112 | | GGGCGGG | 7 | 278 | |
| MR113 | | CATTTAC | 7 | 312 | |
| MR114 | | GACCCAG | 7 | 389 | |
| MR115 | | CTCAACTC | 8 | 421 | |
| MR116 | | ATTTTTA | 7 | 455 | |
| MR117 | | ATTTTTA | 7 | 484 | |
| MR118 | | AGAAAGA | 7 | 526 | 23 |
| MR119 | | AAAGAGAAA | 9 | 528 | |
| MR120 | | AAGGGAA | 7 | 585 | |
| MR121 | | GATATAG | 7 | 629 | |
| MR122 | | GAAAGAAAG | 9 | 657 | |
| MR123 | | AGACCCCAGA | 10 | 790 | |
| MR124 | | CAGTCATACTAAC | 11 | 813 | |
| MR125 | | CCAGTGACC | 9 | 863 | |
| MR126 | | AAGAGGGAGAA | 11 | 892 | |
| MR127 | | ATGACCAGTA | 10 | 927 | |
| MR128 | | GTACATATACATG | 13 | 934 | |
| REGION 7 | | | | | |
| MR129 | | AGGTGGA | 7 | 165 | |
| MR130 | | AGAGGAGA | 8 | 97 | |
| MR131 | | GAGAAGAG | 8 | 132 | |
| MR132 | | AGAGGAGA | 8 | 136 | |
| MR133 | | GCATCAACTACG | 12 | 182 | |
| MR134 | | AGACAGA | 7 | 195 | |
| MR135 | | CAAGAAC | 7 | 222 | |
| MR136 | | AAGTTGAA | 8 | 254 | |
| MR137 | | AAAGAAGGGGAAAGGAAGAAA | 17 | 271 | |
| MR138 | | CTAGGATC | 8 | 314 | |
| MR139 | | AAAGAATTTAAGGAA | 13 | 343 | |
| MR140 | | TTTAAGGAATTT | 12 | 349 | |
| MR141 | | AGTCTCTGA | 9 | 374 | |
| MR142 | | ATGGGTA | 7 | 403 | 25 |
| MR143 | | GACTCAG | 7 | 426 | |
| MR144 | | GCACACG | 7 | 468 | |
| MR145 | | GGCAGGTGGAAGG | 11 | 480 | |
| MR146 | | GGGAGGG | 7 | 570 | |
| MR147 | | CATAATCACGGCTAGCATCCTCCTATGGTACGCACAAATAC | 31 | 636 | |
| MR148 | | CTTATTC | 7 | 730 | |
| MR149 | | AGACAACCAACTGA | 12 | 765 | |
| MR150 | | GCAAACG | 7 | 823 | |
| MR151 | | CTAGAAAAAACGAAGAAAGATC | 18 | 841 | |
| MR152 | | CGAGAGC | 7 | 894 | |
| MR153 | | CTACGTCCTGCATC | 14 | 919 | |
| REGION 8 | | | | | |
| MR154 | | GGAAAGG | 7 | 61 | |
| MR155 | | TTCCCCTT | 8 | 97 | |
| MR156 | | ACTCTCA | 7 | 141 | |

| | | | | | | | |
|---|---|---|---|---|---|---|---|
| MR157 | | TCTTTTCT | | 8 | 155 | | |
| MR158 | | AGAAAAGA | | 8 | 226 | | |
| MR159 | | AACCCAA | | 7 | 255 | | |
| MR160 | | ACCTAGATCCA | | 11 | 286 | | |
| MR161 | | AAAGTTTGAAA | | 11 | 311 | | |
| MR162 | | CCGGGCC | | 7 | 418 | | |
| MR163 | | AGGTTTTGGA | | 10 | 456 | | |
| MR164 | | ATTTTTA | | 7 | 495 | | |
| MR165 | | GAGGGAG | | 7 | 502 | | |
| MR166 | | TTCTCTTTTCTATT | | 12 | 529 | | |
| MR167 | | GAAGAACACAACCAACACAAGAAG | | 24 | 545 | | |
| MR168 | | AAGGGGAA | | 8 | 566 | | |
| MR169 | | GAAAAAG | | 7 | 631 | | |
| MR170 | | AAGTGAA | | 7 | 635 | 26 | |
| MR171 | | AAAGAGGAGAAA | | 12 | 706 | | |
| MR172 | | CGCTGTGTCGC | | 11 | 728 | | |
| MR173 | | AAGGGAA | | 7 | 790 | | |
| MR174 | | AAGGGAA | | 7 | 831 | | |
| MR175 | | AGAATGTAAGA | | 11 | 853 | | |
| MR176 | | AAGAGAA | | 7 | 860 | | |
| MR177 | | AGGAGGA | | 7 | 884 | | |
| MR178 | | AGGACCAGGA | | 10 | 887 | | |
| MR179 | | AGTGTGA | | 7 | 985 | | |
| REGION 9 | | | | | | | |
| MR180 | | AAGCAGGACGAA | | 12 | 39 | | |
| MR181 | | AACAACAA | | 8 | 86 | | |
| MR182 | | ACAACAACA | | 9 | 87 | | |
| MR183 | | TACAAAGGAAATAT | | 12 | 159 | | |
| MR184 | | CACTCTCAC | | 9 | 195 | | |
| MR185 | | GATTGAAAGTGAG | | 11 | 367 | | |
| MR186 | | AGATACCAAACC-TAGA | | 15 | 378 | | |
| MR187 | | AAATAAA | | 7 | 417 | | |
| MR188 | | AAACAAA | | 7 | 498 | | |
| MR189 | | ACAAAACA | | 8 | 500 | | |
| MR190 | | GACACAG | | 7 | 583 | | |
| MR191 | | ACAGACA | | 7 | 599 | | |
| MR192 | | GACAACAG | | 8 | 618 | | |
| MR193 | | AAAGAGAAA | | 9 | 635 | | |
| MR194 | | AGAACCCAAGA | | 11 | 653 | | |
| MR195 | | AACCCAAGAACCGAA | | 13 | 655 | | |
| MR196 | | AAAGAAGGCACG-AAGAAA | | 17 | 668 | | |
| MR197 | | AGAAAAAGA | | 9 | 729 | | 27 |
| MR198 | | GAACAAG | | 7 | 817 | | |
| MR199 | | GGTTTTGG | | 8 | 858 | | |
| MR200 | | AAGGAAAGGAA | | 11 | 878 | | |
| MR201 | | GAAAGGAATCTCCATCTTGAAGGAAAG | | 21 | 881 | | |
| MR202 | | AAAGTGTGAAA | | 11 | 904 | | |
| MR203 | | AACATGTGTGTACAA | | 15 | 913 | | |
| MR204 | | GTACAACATG | | 10 | 922 | | |
| MR205 | | GAAAAAGAGAGAAGAAG | | 15 | 936 | | |
| MR206 | | GTACATG | | 7 | 994 | | |
| REGION 10 | | | | | | | |
| MR207 | | CTCCAGAGAGAACTC | | 13 | 63 | | |
| MR208 | | ATCACACTA | | 9 | 193 | | |
| MR209 | | AAGACCTAAAAAAATGAAGAA | | 16 | 203 | | |
| MR210 | | AATGGTAA | | 8 | 222 | | |
| MR211 | | AGAACACAAGA | | 11 | 246 | | |
| MR212 | | GAGAAGAG | | 8 | 351 | | |
| MR213 | | ACTAATCA | | 8 | 423 | | |
| MR214 | | CTAACAATC | | 9 | 472 | | |
| MR215 | | GTGTTGTG | | 8 | 563 | | |
| MR216 | | ACATTGGGTTCCA | | 11 | 924 | | |
| MR217 | | TCCAACAAGTCGAACAACCT | | 18 | 933 | | |
| MR218 | | ACAAGTCGAACA-ACCTGGTCCATACATGCTAAACA | | 30 | 937 | | |
| MR219 | | CTGACAGTC | | 9 | 997 | | |

| | | | | | |
|---|---|---|---|---|---|
| MR220 | | ATTGGGTTA | 9 | 1120 | |
| MR221 | | AGAAGAGAAGA | 11 | 1237 | 23 |
| MR222 | | AGAAG-AGAAGAGGAAGA | 16 | 1237 | |
| MR223 | | GAAGAGGAAGAAGCAGGAG | 15 | 1243 | |
| MR224 | | TAAAAAAT | 8 | 1443 | |
| MR225 | | GAAGCTGTACGCATGGCGTAG | 17 | 1471 | |
| MR226 | | AGAGGAGA | 8 | 1505 | |
| MR227 | | AGAGGAGA | 8 | 1592 | |
| MR228 | | ACAAAAAACA | 10 | 1610 | |
| MR229 | | AGACCAGA | 8 | 1638 | |
| | | RABIES VIRUS | | | |
| REGION 1 | | | | | |
| MIRROR REPEATS | | SEQUENCE | LENGTH | POSITION IN REGION | TOTAL |
| MR1 | | AAAGAAA | 7 | 19 | |
| MR2 | | ACAGACA | 7 | 28 | |
| MR3 | | AATGTAA | 7 | 54 | |
| MR4 | | ATCAATATGAGTACAAGTA | 15 | 141 | |
| MR5 | | GAAAAAG | 7 | 178 | |
| MR6 | | CTAGGAAAGGCTC | 11 | 197 | |
| MR7 | | ATAAAAA | 7 | 217 | |
| MR8 | | TGTATGT | 7 | 277 | |
| MR9 | | GGCGGCGG | 8 | 292 | |
| MR10 | | CGGCGGC | 7 | 294 | |
| MR11 | | AGTTTTTTGA | 10 | 306 | |
| MR12 | | GATGTAG | 7 | 413 | |
| MR13 | | AAGGGAA | 7 | 420 | |
| MR14 | | GTCCCTG | 7 | 470 | |
| MR15 | | TTCTCTT | 7 | 498 | 24 |
| MR16 | | ACAAACA | 7 | 557 | |
| MR17 | | GTCACTG | 7 | 758 | |
| MR18 | | TGTTCAGGACTGGT | 12 | 776 | |
| MR19 | | ACTGGTGTCATTTACTGGGTTCA | 19 | 784 | |
| MR20 | | AAACAAA | 7 | 809 | |
| MR21 | | CAAGAAC | 7 | 856 | |
| MR22 | | TAAGAAGAAT | 10 | 876 | |
| MR23 | | CTCACTC | 7 | 915 | |
| MR24 | | AATCTCCTTATTCATCAAA | 15 | 960 | |
| REGION 2 | | | | | |
| MR25 | | TTCACTT | 7 | 5 | |
| MR26 | | GTAGGATG | 8 | 13 | |
| MR27 | | ATCCCTA | 7 | 39 | |
| MR28 | | GTTATTG | 7 | 55 | |
| MR29 | | TGTCTGT | 7 | 83 | |
| MR30 | | GGAGAGG | 7 | 106 | |
| MR31 | | GGGAAAGGG | 9 | 121 | |
| MR32 | | AGAAAGA | 7 | 158 | |
| MR33 | | AAGAACTTCAAGAA | 14 | 161 | |
| MR34 | | AACTGTCAA | 9 | 225 | |
| MR35 | | TGGAGGT | 7 | 306 | |
| MR36 | | AAGAAGTTGAATAA | 12 | 422 | |
| MR37 | | AGCCGGTCTGGCCGA | 15 | 549 | |
| MR38 | | CCAAGGGGAACC | 12 | 630 | |
| MR39 | | CCCAACCC | 8 | 703 | 24 |
| MR40 | | GACTTTCAG | 9 | 751 | |
| MR41 | | AGGAGAGGA | 6 | 768 | |
| MR42 | | GATCCTAG | 8 | 775 | |
| MR43 | | GAGAGAG | 7 | 845 | |
| MR44 | | AGAGAGA | 7 | 846 | |
| MR45 | | TATCCTAT | 8 | 890 | |
| MR46 | | TCCCAACCCT | 10 | 912 | |
| MR47 | | CGAGAGC | 7 | 964 | |
| MR48 | | CACCCAC | 7 | 986 | |
| REGION 3 | | | | | |

| | | | | | | |
|---|---|---|---|---|---|---|
| MR49 | | AGAGAGA | | 1 | 7 | |
| MR50 | | CGAAAGC | | 7 | 19 | |
| MR51 | | TCAAACT | | 7 | 38 | |
| MR52 | | AATATAA | | 7 | 148 | |
| MR53 | | TGATATAGT | | 9 | 218 | |
| MR54 | | TCCCCCT | | 7 | 283 | |
| MR55 | | ATGTGTA | | 7 | 293 | |
| MR56 | | AGTCGAATCCAAC--AAGCTGA | | 18 | 344 | |
| MR57 | | ATCGCTA | | 7 | 388 | |
| MR58 | | ACCTCTCCA | | 9 | 412 | |
| MR59 | | GCAGGGACG | | 9 | 527 | |
| MR60 | | CAAAAAC | | 7 | 544 | |
| MR61 | | CCCTCTCCC | | 9 | 550 | 21 |
| MR62 | | CAAGAAGAAC | | 10 | 633 | |
| MR63 | | AGGAGGA | | 7 | 643 | |
| MR64 | | AGCCCGA | | 7 | 682 | |
| MR65 | | GTTATTG | | 7 | 781 | |
| MR66 | | AGTCCCTGA | | 9 | 810 | |
| MR67 | | ACTCTCA | | 7 | 908 | |
| MR68 | | GAGTGAG | | 7 | 962 | |
| MR69 | | GTGAGTG | | 7 | 964 | |
| REGION 4 | | | | | | |
| MR70 | | GTATCAACATGAACTCGAGAGCAGGT-CAACTATG | | 27 | 4 | |
| MR71 | | CTCAACTC | | 8 | 163 | |
| MR72 | | CAATATAAC | | 9 | 184 | |
| MR73 | | TTTACATTT | | 9 | 253 | |
| MR74 | | TGTTTGT | | 7 | 337 | |
| MR75 | | TTGTGTT | | 7 | 363 | |
| MR76 | | GGTCCCTGG | | 9 | 408 | |
| MR77 | | AGCCCGA | | 7 | 417 | |
| MR78 | | ACATACA | | 7 | 427 | |
| MR79 | | TAAAAAT | | 7 | 535 | |
| MR80 | | GTTCACTTG | | 9 | 548 | 19 |
| MR81 | | CGGAGGC | | 7 | 571 | |
| MR82 | | AAAGAAA | | 7 | 622 | |
| MR83 | | TGGCCGGT | | 8 | 679 | |
| MR84 | | GACCCCAG | | 8 | 687 | |
| MR85 | | AACCACCAA | | 9 | 752 | |
| MR86 | | TAGGGAT | | 7 | 928 | |
| MR87 | | CATTTTTAC | | 9 | 944 | |
| MR88 | | GAGTGAG | | 7 | 983 | |
| REGION 5 | | | | | | |
| MR89 | | GTAGATG | | 7 | 2 | |
| MR90 | | AACTCAA | | 7 | 45 | |
| MR91 | | AACTCAA | | 8 | 117 | |
| MR92 | | GCCCTCCCG | | 9 | 129 | |
| MR93 | | AGAAGAGAGAGGA | | 11 | 207 | |
| MR94 | | GAAGAGAGAGGAG | | 11 | 208 | |
| MR95 | | ACCACCA | | 7 | 248 | |
| MR96 | | GTCCCTG | | 7 | 296 | |
| MR97 | | GGTTTGG | | 7 | 303 | |
| MR98 | | GAAAAGCATATACCATATTCAACAAG | | 20 | 309 | |
| MR99 | | TATAATAT | | 8 | 460 | |
| MR100 | | CCTCCTCC | | 8 | 511 | |
| MR101 | | GTTGTTG | | 7 | 532 | |
| MR102 | | TCCCCCT | | 7 | 552 | |
| MR103 | | CAAGAAC | | 7 | 592 | |
| MR104 | | TTGAAGTT | | 8 | 621 | |
| MR105 | | TATGTAT | | 7 | 692 | |
| MR106 | | GAGAAGAG | | 8 | 757 | 26 |
| MR107 | | ACACAACACA | | 10 | 782 | |
| MR108 | | AACACAA | | 7 | 786 | |
| MR109 | | GAGGGACAGGGAG | | 13 | 798 | |
| MR110 | | AGGGAGGGA | | 9 | 805 | |

| | | | | | | |
|---|---|---|---|---|---|---|
| MR111 | | GGAGGGAGG | | 9 | 807 | |
| MR112 | | AAGTCCTGAA | | 10 | 918 | |
| MR113 | | CTTGGGGGGTTC | | 12 | 939 | |
| MR114 | | TCCTCCT | | 7 | 979 | |
| REGION 6 | | | | | | |
| MR115 | | GATCTTCTAG | | 10 | 83 | |
| MR116 | | TCAGTGACT | | 9 | 98 | |
| MR117 | | ACAGACA | | 7 | 189 | |
| MR118 | | GGACTCAGG | | 9 | 222 | |
| MR119 | | GAGTTAATTGAG | | 12 | 233 | 20 |
| MR120 | | GAGAGAG | | 7 | 242 | |
| MR121 | | CTGGGTC | | 7 | 330 | |
| MR122 | | CTGGAGAGGTC | | 11 | 428 | |
| MR123 | | GACCCAA-TCG-AGTTAGAGGCTGAACCCAG | | 25 | 454 | |
| MR124 | | ACCCCCA | | 7 | 487 | |
| MR125 | | TCTCAACTCT | | 10 | 528 | |
| MR126 | | TAGAAGAT | | 8 | 545 | |
| MR127 | | TAACAGACAAT | | 11 | 611 | |
| MR128 | | AGTTTTGA | | 8 | 639 | |
| MR129 | | TTGGGTT | | 7 | 670 | |
| MR130 | | TCTCTCT | | 7 | 715 | |
| MR131 | | GAGGGAG | | 7 | 859 | |
| MR132 | | TGGAAGGT | | 8 | 906 | |
| MR133 | | CTTGTTC | | 7 | 939 | |
| MR134 | | AGAAAAGA | | 8 | 990 | |
| REGION 7 | | | | | | |
| MR135 | | TTTGTTT | | 7 | 99 | |
| MR136 | | TCTTTTCT | | 8 | 161 | |
| MR137 | | CAGAGCCCCGATAC | | 12 | 208 | |
| MR138 | | TATTTAT | | 7 | 400 | |
| MR139 | | ACATACA | | 7 | 487 | |
| MR140 | | GTTTTTG | | 7 | 501 | |
| MR141 | | ATATATA | | 7 | 539 | |
| MR142 | | TCAAAACT | | 8 | 564 | |
| MR143 | | ATAGATA | | 7 | 600 | |
| MR144 | | AGGAGGA | | 7 | 642 | 14 |
| MR145 | | AGCCCGA | | 7 | 704 | |
| MR146 | | AAACCCAAA | | 9 | 736 | |
| MR147 | | GAAAAAG | | 7 | 930 | |
| MR148 | | TCTGAAGTCT | | 10 | 983 | |
| REGION 8 | | | | | | |
| MR149 | | GAATCTAAG | | 9 | 94 | |
| MR150 | | CTACATC | | 7 | 139 | |
| MR151 | | ACAGACA | | 7 | 173 | |
| MR152 | | GAACAAG | | 7 | 184 | |
| MR153 | | TGAAAAGT | | 8 | 274 | |
| MR154 | | AGATTAGA | | 8 | 296 | |
| MR155 | | TTTCTGTCCTAGATCAAGTGTTT | | 17 | 321 | |
| MR156 | | AGACAGA | | 7 | 406 | |
| MR157 | | GTCCAACGGCCCAACCTG | | 14 | 460 | |
| MR158 | | GATAGATAG | | 9 | 547 | |
| MR159 | | AGATAGA | | 7 | 550 | 20 |
| MR160 | | AAATCAGGAACACAAGAACCAAA | | 19 | 564 | |
| MR161 | | ATCAGGAACACAAGAACCAAAGTACTA | | 21 | 566 | |
| MR162 | | AAGAAAGAA | | 9 | 749 | |
| MR163 | | ATGTGTA | | 7 | 764 | |
| MR164 | | TTTGTTT | | 7 | 802 | |
| MR165 | | CCTGAGTCC | | 9 | 827 | |
| MR166 | | AATATAA | | 7 | 890 | |
| MR167 | | CACAACAC | | 8 | 930 | |
| MR168 | | ACTCTCA | | 7 | 936 | |
| REGION 9 | | | | | | |

| | | | | |
|---|---|---|---|---|
| MR169 | TCGACCCAGCT | 11 | 934 | 1 |
| REGION 10 | | | | |
| MR170 | AAACT-GGTTCATTACTAGAGATTCCAACTTGGCTCAAA | 29 | 14 | |
| MR171 | CTCAAACTC | 9 | 46 | |
| MR172 | GTCTCTG | 7 | 71 | |
| MR173 | CTCTCTC | 7 | 198 | |
| MR174 | ACAGACA | 7 | 219 | |
| MR175 | ACAGACA | 7 | 293 | |
| MR176 | ACAGAGAGACA | 11 | 317 | 15 |
| MR177 | ACCTCCA | 7 | 358 | |
| MR178 | GTGTGTG | 7 | 374 | |
| MR179 | GAGATTTTGAATCTCTAAGCGGTAGAG | 21 | 514 | |
| MR180 | GACTCAG | 7 | 603 | |
| MR181 | CATATAC | 7 | 641 | |
| MR182 | TCTTTTAGAGAAGAGATATTTTCT | 20 | 840 | |
| MR183 | TTGTGTT | 7 | 924 | |
| MR184 | GAGCGAG | 7 | 954 | |
| REGION 11 | | | | |
| MR185 | TTTCAGACTTT | 11 | 3 | |
| MR186 | TCATTACT | 8 | 42 | |
| MR187 | CTCATCTTCTACTC | 14 | 57 | |
| MR188 | AGAGTATGAGA | 11 | 96 | |
| MR189 | AGTTCCTTGA | 10 | 125 | |
| MR190 | GTAGGATG | 8 | 305 | |
| MR191 | CCCGGCCC | 8 | 364 | |
| MR192 | CCATCTCTCTGCC | 11 | 506 | |
| MR193 | AGGGGGGA | 8 | 538 | |
| MR194 | CTTGTGTTC | 9 | 584 | |
| MR195 | GAGGGGAG | 8 | 661 | 20 |
| MR196 | GGTCTTCAAAACTTATGG | 16 | 895 | |
| MR197 | GGTCTTCAAAACTTATGG | 9 | 1128 | |
| MR198 | GTTCCTTG | 8 | 1149 | |
| MR199 | ATCTTTTCTA | 10 | 1249 | |
| MR200 | GGGGAACTCTGTCTAAAGTGG | 17 | 1293 | |
| MR201 | ATGTATCTATCTA | 11 | 1448 | |
| MR202 | ATCTATCTA | 9 | 1452 | |
| MR203 | AGATCTAGA | 9 | 1763 | |
| MR204 | AGATCATCCCTACTAGA | 17 | 1769 | |

HTLV- 1

REGION 1

| MIRROR REPEATS | SEQUENCE | LENGTH | POSITION IN REGION | TOTAL |
|---|---|---|---|---|
| MR1 | CGCCCGC | 7 | 92 | |
| MR2 | CCGCCGCC | 8 | 95 | |
| MR3 | CCGGGCC | 7 | 143 | |
| MR4 | GACTCAG | 7 | 180 | |
| MR5 | CTCAACTC | 8 | 219 | |
| MR6 | CTTTGTTTC | 9 | 232 | |
| MR7 | CCCTTTCCC | 9 | 279 | |
| MR8 | CTTTCCCTTTC | 11 | 281 | |
| MR9 | ACGGCCAAGTACCGGCA | 17 | 319 | |
| MR10 | GGCTCGG | 7 | 344 | |
| MR11 | CAGCGAC | 7 | 354 | |
| MR12 | CGACAGC | 7 | 357 | 24 |
| MR13 | CGGGGGC | 7 | 498 | |
| MR14 | GGTCTGG | 7 | 620 | |
| MR15 | CTCCCTC | 7 | 644 | |
| MR16 | TACACAT | 7 | 700 | |
| MR17 | ACATACTCATCCA | 13 | 703 | |
| MR18 | CCAAACCCAAGCC | 13 | 713 | |
| MR19 | CACCCAC | 7 | 767 | |
| MR20 | TCCCCCCT | 8 | 799 | |
| MR21 | CCCTCCC | 7 | 803 | |
| MR22 | CCTTCCAGTCATGCACCCACATGGTGCCCCTCC | 33 | 836 | |
| MR23 | CCCTCCC | 7 | 863 | |

| | | | | | |
|---|---|---|---|---|---|
| MR24 | | CAGTTTGAC | 9 | 978 | |
| REGION 2 | | | | | |
| MR25 | | CCTTTGCTCCTCCCTCGTGGCTTCC | 25 | 22 | |
| MR26 | | CCTCCCT-CGTGGCTTCCCTCC | 22 | 30 | |
| MR27 | | ACAACAACA | 9 | 151 | |
| MR28 | | AACAACAA | 8 | 153 | |
| MR29 | | CATTTTAC | 8 | 340 | |
| MR30 | | AAACAAA | 7 | 370 | |
| MR31 | | ACCCCCA | 7 | 455 | |
| MR32 | | CCCCCAAAGACAAAACCAAAGTGTTAGTTGTCCAGCCTAAA---AA | 52 | 456 | |
| MR33 | | TCCCCCT | 7 | 574 | |
| MR34 | | GCCCCCG | 7 | 627 | |
| MR35 | | AGGAGGA | 7 | 666 | |
| MR36 | | CCCACACCC | 9 | 703 | |
| MR37 | | CAAAAAAC | 8 | 711 | |
| MR38 | | ACATTACA | 8 | 751 | |
| MR39 | | TCCTTCCT | 8 | 764 | 25 |
| MR40 | | CCCGCCC | 7 | 814 | |
| MR41 | | CCAGACC | 7 | 852 | |
| MR42 | | TCGAAGCT | 8 | 875 | |
| MR43 | | TGACAGT | 7 | 905 | |
| MR44 | | TTCCGATAGCCTT | 13 | 914 | |
| MR45 | | CTTGTTC | 7 | 924 | |
| MR46 | | AATACTCCCCTCAAAA | 16 | 937 | |
| MR47 | | CAAAAAC | 7 | 948 | |
| MR48 | | CCAAACC | 7 | 978 | |
| MR49 | | AACCCAA | 7 | 981 | |
| REGION 3 | | | | | |
| MR50 | | CCTTTCC | 7 | 30 | |
| MR51 | | TAGTTGAT | 8 | 67 | |
| MR52 | | TAGTTGAT | 8 | 77 | |
| MR53 | | AAGGCCCCCCGAAA | 14 | 212 | |
| MR54 | | TTCCCTT | 7 | 234 | |
| MR55 | | CCGGAAGGCC | 10 | 277 | |
| MR56 | | ACCGGGCCA | 9 | 314 | |
| MR57 | | ACTCTCTA---ACCATAGATCTCTCA | 26 | 396 | |
| MR58 | | TCCCCCGGGCCCCCT | 15 | 425 | |
| MR59 | | CCTTTTTCC | 9 | 498 | |
| MR60 | | CAAATCCCCCTACCTAAAC | 19 | 506 | 19 |
| MR61 | | ACGGCCCCGGCA | 12 | 570 | |
| MR62 | | TAAAAATAGTCCCACCCTGTTCGAAAT | 27 | 616 | |
| MR63 | | GTACATG | 7 | 703 | |
| MR64 | | CCCTCCC | 7 | 731 | |
| MR65 | | ACTCTCA | 7 | 754 | |
| MR66 | | GCCTGTGTCCG | 11 | 796 | |
| MR67 | | CCGAAAACAAAACCC | 15 | 804 | |
| MR68 | | CCTGAACTTCAAGCCC | 16 | 923 | |
| REGION 4 | | | | | |
| MR69 | | CCAAACC | 7 | 123 | |
| MR70 | | TGGCACCACCACTGT | 15 | 165 | |
| MR71 | | TGTGGTGT | 8 | 177 | |
| MR72 | | CCAAACC | 7 | 324 | |
| MR73 | | CCAAACC | 7 | 351 | |
| MR74 | | TCAGACGGA---TCCACCTCCCAGGCAGCCT | 31 | 562 | |
| MR75 | | TTCCCCCTT | 9 | 628 | |
| MR76 | | CGCCACCGC | 9 | 638 | 15 |
| MR77 | | CCTCTCC | 7 | 690 | |
| MR78 | | GGCACCTTCCAAGG | 14 | 775 | |
| MR79 | | TTGCACCACGTT | 12 | 853 | |

| | | | | | |
|---|---|---|---|---|---|
| MR80 | | CCATACC | 7 | 870 | |
| MR81 | | CCCTACTAATCACCCC | 16 | 920 | |
| MR82 | | CCTGTCC | 7 | 934 | |
| MR83 | | CTCTCTC | 7 | 946 | |
| REGION 5 | | | | | |
| MR84 | | ACATTACCCATTTCA | 15 | 136 | |
| MR85 | | AAATATAAA | 9 | 150 | |
| MR86 | | ATAAAAATA | 9 | 154 | |
| MR87 | | ATGGGTA | 7 | 182 | |
| MR88 | | AAAAGAGAAAA | 11 | 220 | |
| MR89 | | AGAAAAGA | 8 | 225 | |
| MR90 | | AAAGAAA | 7 | 228 | |
| MR91 | | ACAGACA | 7 | 303 | |
| MR92 | | ATGTGTA | 7 | 345 | |
| MR93 | | TACCCAT | 7 | 374 | |
| MR94 | | ATTCTTA | 7 | 429 | |
| MR95 | | TCACCACT | 8 | 563 | |
| MR96 | | CAGAGAC | 7 | 592 | 21 |
| MR97 | | AAACAAA | 7 | 615 | |
| MR98 | | GGAAAGG | 7 | 667 | |
| MR99 | | CGGAGGC | 7 | 788 | |
| MR100 | | AAAGAAA | 7 | 807 | |
| MR101 | | AGAAAAAGA | 9 | 809 | |
| MR102 | | ACCACCA | 7 | 817 | |
| MR103 | | ACCACCA | 7 | 829 | |
| MR104 | | ACTCTCA | 7 | 910 | |
| REGION 6 | | | | | |
| MR105 | | TATTCCCTATATCTAT-TCCCTCAT | 25 | 66 | |
| MR106 | | AAAGCCAAACCGAAA | 15 | 98 | |
| MR107 | | CGGAGGC | 7 | 116 | |
| MR108 | | CTTATTC | 7 | 136 | |
| MR109 | | CCTTGTTCC | 9 | 147 | |
| MR110 | | CCATACC | 7 | 165 | |
| MR111 | | ACCATATC-CTCGAGCCCTCTATACCA | 27 | 418 | |
| MR112 | | TCAAAACT | 8 | 450 | |
| MR113 | | TGTCTGT | 7 | 506 | |
| MR114 | | CCTATCC | 7 | 527 | |
| MR115 | | GTCTCTG | 7 | 561 | |
| MR116 | | CTTCTTC | 7 | 577 | |
| MR117 | | TACCCAT | 7 | 597 | 25 |
| MR118 | | TTACCATT | 8 | 633 | |
| MR119 | | CTTTGACCCCCAGATTC | 17 | 656 | |
| MR120 | | TCCCCCTGTCATAACTCCCTC-ATCCTGCCCCCCT | 35 | 687 | |
| MR121 | | CCTTTTCC | 8 | 718 | |
| MR122 | | CCTTGTCACCTGTTCC | 16 | 724 | |
| MR123 | | CTAGGATC | 8 | 744 | |
| MR124 | | CGGTGGC | 7 | 772 | |
| MR125 | | TGGCGGT | 7 | 775 | |
| MR126 | | CGGTCTGGC | 9 | 778 | |
| MR127 | | AGGTGGA | 7 | 874 | |
| MR128 | | AACTCAA | 7 | 899 | |
| MR129 | | CAAAAAC | 7 | 914 | |
| REGION 7 | | | | | |
| MR130 | | ACAAGAACA | 9 | 16 | |
| MR131 | | AGAGAGA | 7 | 70 | |
| MR132 | | AGTCCTGA | 8 | 94 | |
| MR133 | | TCGCGCT | 7 | 177 | |
| MR134 | | CTCCCCTC | 8 | 236 | |

| | | | | | |
|---|---|---|---|---|---|
| MR135 | | TACCCCCAT | 9 | 254 | |
| MR136 | | CTGAGTC | 7 | 279 | |
| MR137 | | AATTAACCTCTCCCATCAA | 19 | 347 | |
| MR138 | | TCCTCCT | 7 | 367 | |
| MR139 | | CAGCAACCTCCTCCGTTCAGCCTCCAAGGAC | 31 | 381 | |
| MR140 | | CTCCACCTC | 9 | 411 | |
| MR141 | | TCAACCCCCAACT | 13 | 445 | |
| MR142 | | TTTCTTT | 7 | 467 | |
| MR143 |  | CCTTCTCCTCCGCC----CGCTTCCTGCGCCGTGCCTTCTCCTCTT | 48 | 569 | 27 |
| MR144 | | CCTTCTCCTCTTCC | 14 | 599 | |
| MR145 | | CTCTTCCTTCCTTTTC | 16 | 606 | |
| MR146 | | TTTTCCTCCTCTT | 13 | 640 | |
| MR147 | | CTCCTCTTTCTCCCGCTCttttttttCGCTTCCTCTT-CTCCTC | 43 | 645 | |
| MR148 | | CCGTCGCTGCC | 11 | 690 | |
| MR149 | | TCCCCCCT | 8 | 734 | |
| MR150 | | GCCGGCCG | 8 | 758 | |
| MR151 | | TAGAGAT | 7 | 780 | |
| MR152 | | CAGTTTCCTCCTCCTCCTTGTCCTTTAAC | 29 | 803 | |
| MR153 | | TTCCTCCTCCTCCTT | 15 | 807 | |
| MR154 | | CCTCCTTGTCCTTTAACTCTTCCTCC | 26 | 815 | |
| MR155 | | GATAATAG | 8 | 844 | |
| MR156 | | GGTTTGG | 7 | 963 | |
| REGION 8 | | | | | |
| MR157 | | ACCTGTCCA | 9 | 88 | |
| MR158 | | CTCCCCTC | 8 | 169 | |
| MR159 | | TACAACCCCCAACAT | 15 | 234 | |
| MR160 | | CCTCCTTCCTCC | 12 | 254 | |
| MR161 | | TCCCAACCCT | 10 | 329 | |
| MR162 | | ACCCCGGACTCCGGCCCCA | 19 | 350 | |
| MR163 | | CCAAAACC | 8 | 366 | |
| MR164 | | TCTGCATGTACCTCT | 15 | 401 | |
| MR165 | | TTTGCCCACCACCCTTTT | 18 | 582 | |
| MR166 | | CCTTTTCC | 8 | 594 | |
| MR167 | | GGCAAAACGG | 10 | 635 | |
| MR168 | | CCTCACCACTCC | 12 | 666 | |
| MR169 | | TCCTCCTCCT | 10 | 763 | 19 |
| MR170 | | CTTTATATTTC | 11 | 771 | |
| MR171 | | TCAAAAAACT | 10 | 1201 | |
| MR172 | | ACTTTTCA | 8 | 1208 | |
| MR173 | | CAATAAAATAAC | 12 | 1232 | |
| MR174 | | CCGCCGCC | 8 | 1304 | |
| MR175 | | CTCAACTC | 8 | 1498 | |
| | | TMV | | | |
| REGION 1 | | | | | |
| MIRROR REPEATS | | SEQUENCE | LENGTH | POSITION IN REGION | TOTAL |
| MR1 | | AATGATCTAGCAA | 11 | 138 | |
| MR2 | | CGACACAGC | 9 | 164 | |
| MR3 | | AACTTTTCAA | 10 | 213 | |
| MR4 | | CGAGGAGC | 8 | 233 | |
| MR5 | | GCAGACG | 7 | 239 | |
| MR6 | | TTACATT | 7 | 280 | |
| MR7 | | TTCGCTT | 7 | 311 | |
| MR8 | | AGGTGGA | 7 | 320 | |
| MR9 | | ACGAGCA | 7 | 425 | 17 |
| MR10 | | CAGAAAGAC | 9 | 489 | |
| MR11 | | AGAGAGA | 7 | 524 | |
| MR12 | | GAGAGAG | 7 | 525 | |
| MR13 | | AGGGGGGA | 8 | 530 | |
| MR14 | | CGCAGAAATTCCTGAAGACGC | 17 | 578 | |
| MR15 | | ACGCTGTCTGTCACA | 13 | 595 | |

| | | | | | |
|---|---|---|---|---|---|
| MR16 | | ACCTTTTCTTTTGCA | 13 | 843 | |
| MR17 | | ATTCTTA | 7 | 897 | |
| REGION 2 | | | | | |
| MR18 | | ATTCTTA | 7 | 5 | |
| MR19 | | TTTTCTTTT | 9 | 13 | |
| MR20 | | GGTGTGG | 7 | 29 | |
| MR21 | | TCCTCCT | 7 | 129 | |
| MR22 | | TTTCTTT | 7 | 207 | |
| MR23 | | ACGCGCA | 7 | 230 | |
| MR24 | | CGAAAGC | 7 | 300 | 11 |
| MR25 | | TGTTTTGT | 8 | 322 | |
| MR26 | | AAGCTTGCCGTTCTAA | 14 | 446 | |
| MR27 | | TAGAGAT | 7 | 615 | |
| MR28 | | ACCTTCCA | 8 | 644 | |
| REGION 3 | | | | | |
| MR29 | | AGAGGAGA | 8 | 16 | |
| MR30 | | GAGATAGAG | 9 | 82 | |
| MR31 | | GGCAACGG | 8 | 111 | |
| MR32 | | AGCTCGA | 7 | 145 | |
| MR33 | | CGATAGC | 7 | 198 | |
| MR34 | | ATCACTA | 7 | 213 | |
| MR35 | | GTTGTTG | 7 | 316 | |
| MR36 | | GGTGTGG | 7 | 418 | |
| MR37 | | GAGAAGAG | 8 | 441 | 19 |
| MR38 | | TCTGAGTCT | 9 | 460 | |
| MR39 | | TGTTCTTGT | 9 | 552 | |
| MR40 | | AAACCAAA | 8 | 584 | |
| MR41 | | AAAGAAA | 7 | 589 | |
| MR42 | | CAGAAGAC | 8 | 663 | |
| MR43 | | GCACACG | 7 | 752 | |
| MR44 | | TTGTGTT | 7 | 810 | |
| MR45 | | GACACACAG | 9 | 865 | |
| MR46 | | CCCGCCC | 7 | 913 | |
| MR47 | | GAGGTGGAG | 9 | 943 | |
| REGION 4 | | | | | |
| MR48 | | CTTCTTC | 7 | 31 | |
| MR49 | | GCCGCCG | 7 | 75 | |
| MR50 | | TCACACT | 7 | 176 | |
| MR51 | | TAGAGAT | 7 | 353 | |
| MR52 | | AGAGATCTAGAGA | 13 | 354 | |
| MR53 | | ATATGTATA | 9 | 388 | |
| MR54 | | AACACAA | 7 | 410 | |
| MR55 | | ACGGCGGCA | 9 | 672 | 16 |
| MR56 | | GGAAAATTTAGTGGCGATGATT-AAAAGG | 25 | 707 | |
| MR57 | | AGTTTTTTGA | 10 | 802 | |
| MR58 | | GTTTTTTGATAGTTATTTG | 17 | 803 | |
| MR59 | | TGTTTCTTTGT | 11 | 851 | |
| MR60 | | TCTCTCA | 7 | 873 | |
| MR61 | | TAACAAT | 7 | 907 | |
| MR62 | | ACAGACA | 7 | 964 | |
| MR63 | | AACCCAA | 7 | 985 | |
| REGION 5 | | | | | |
| MR64 | | TTTGCAGACGATT | 11 | 31 | |
| MR65 | | TTTTTGTTTTT | 11 | 134 | |
| MR66 | | AGAAAGA | 7 | 149 | |
| MR67 | | TTGGGTT | 7 | 293 | |
| MR68 | | AGAAAGA | 7 | 338 | |
| MR69 | | AAAGAAA | 7 | 393 | |
| MR70 | | AGAAAGA | 7 | 395 | 14 |

| | | | | | |
|---|---|---|---|---|---|
| MR71 | | ATCCCCTA | 8 | 666 | |
| MR72 | | ACACACA | 7 | 783 | |
| MR73 | | GTTTGTTTATAAAA--GTCTGGTGAAGTATTTGTCTG | 27 | 838 | |
| MR74 | | TATTTGTCTGATAAAGTTCTTTTTAGA-AGTTTGTTTAT | 29 | 863 | |
| MR75 | | TAGAAGTTTGTTTATAGAT | 15 | 886 | |
| MR76 | | AAAGGAAA | 8 | 918 | |
| MR77 | | AAGTGAA | 7 | 925 | |
| REGION 6 | | | | | |
| MR78 | | ATAAAATA | 8 | 21 | |
| MR79 | | GCCGGTTTGGTCG | 11 | 110 | |
| MR80 | | AGAAAAGA | 8 | 243 | |
| MR81 | | GAAAAAAG | 8 | 546 | |
| MR82 | | AAAGGGAAA | 9 | 566 | |
| MR83 | | TAGTAATGAT | 10 | 580 | |
| MR84 | | TAATTTAAT | 9 | 655 | |
| MR85 | | TCGGAGGCT | 9 | 674 | |
| MR86 | | TGTTCTTGT | 9 | 746 | 20 |
| MR87 | | TTAATTAATT | 10 | 781 | |
| MR88 | | CAAGCTCGAAC | 11 | 829 | |
| MR89 | | AGGTGTGGA | 9 | 863 | |
| MR90 | | TAGAAGTTGAAAAT | 12 | 995 | |
| MR91 | | AGATGCTACTCGTAGA | 14 | 1038 | |
| MR92 | | ATAAGGAGCGCGATAAATA | 15 | 1075 | |
| MR93 | | ATAATTTAATA | 11 | 1091 | |
| MR94 | | TCTT-TCGAGAGCTCTTCT | 17 | 1141 | |
| MR95 | | TCTTCTGGTTTGGTTTGGACCTCT | 20 | 1153 | |
| MR96 | | AATAAATAA | 9 | 1208 | |
| MR97 | | CGTGGTGC | 8 | 1238 | |
| | | PVY | | | |
| REGION 1 | | | | | |
| MIRROR REPEATS | | SEQUENCE | LENGTH | POSITION IN REGION | TOTAL |
| MR1 | | AACTCAA | 7 | 10 | |
| MR2 | | AATACAACATAA | 12 | 15 | |
| MR3 | | AACAACGCAAAAA | 11 | 31 | |
| MR4 | | CAAAAAC | 7 | 38 | |
| MR5 | | CTCACTC | 7 | 62 | |
| MR6 | | CAACTCAAC | 9 | 138 | |
| MR7 | | AACACAA | 7 | 316 | |
| MR8 | | TCGTGTGCT | 9 | 361 | |
| MR9 | | CAAAAAC | 7 | 370 | |
| MR10 | | GGAGAGGAAGGATAGG | 14 | 436 | 23 |
| MR11 | | AGTCTGA | 7 | 528 | |
| MR12 | | ATGCGTA | 7 | 575 | |
| MR13 | | GGAGAAAAGAGG | 12 | 661 | |
| MR14 | | AATATAA | 7 | 711 | |
| MR15 | | TGCAACGT | 8 | 819 | |
| MR16 | | CTCGCTC | 7 | 827 | |
| MR17 | | ACTATCA | 7 | 863 | |
| MR18 | | GGTGATAGTGG | 11 | 881 | |
| MR19 | | AACACAA | 7 | 902 | |
| MR20 | | CTTGTTC | 7 | 946 | |
| MR21 | | GTGCGTG | 7 | 956 | |
| MR22 | | AAGGGAA | 7 | 992 | |
| MR23 | | TGCACGT | 7 | 988 | |
| REGION 2 | | | | | |
| MR24 | | AACTCAA | 7 | 19 | |
| MR25 | | GGTCTGG | 7 | 61 | |
| MR26 | | ATGTGTA | 7 | 108 | |
| MR27 | | TAGCGAT | 7 | 231 | |
| MR28 | | AGTTCTTGA | 9 | 320 | |
| MR29 | | TCGAGCT | 7 | 368 | |

| | | | | | | |
|---|---|---|---|---|---|---|
| MR30 | | AATTTAA | | 7 | 427 | |
| MR31 | | AAAGGAAA | | 8 | 457 | |
| MR32 | | AAAAGAAAA | | 9 | 462 | |
| MR33 | | AGCTCAATTGAGTTTGCTCGA | | 17 | 492 | 21 |
| MR34 | | TCTTTCT | | 7 | 565 | |
| MR35 | | AAATAAA | | 7 | 576 | |
| MR36 | | TAAAAAT | | 7 | 636 | |
| MR37 | | TTTCT-CAAACTTCTTT | | 15 | 687 | |
| MR38 | | GGAACAAGG | | 9 | 760 | |
| MR39 | | GATTTAG | | 7 | 802 | |
| MR40 | | TATGTGTAT | | 9 | 892 | |
| MR41 | | GTTGTTG | | 7 | 905 | |
| MR42 | | CACAACAC | | 8 | 912 | |
| MR43 | | AATCAACATTCTATCCACCAACTAA | | 19 | 941 | |
| MR44 | | AGTGGTGA | | 8 | 988 | |
| REGION 3 | | | | | | |
| MR45 | | GTTATTG | | 7 | 55 | |
| MR46 | | TATTGTTAT | | 9 | 57 | |
| MR47 | | TTATATT | | 7 | 62 | |
| MR48 | | AGGAGGA | | 7 | 103 | |
| MR49 | | AAAGAAA | | 7 | 125 | |
| MR50 | | TGTGTGT | | 7 | 142 | |
| MR51 | | GTGTGTG | | 7 | 143 | |
| MR52 | | GTACATG | | 7 | 225 | |
| MR53 | | AGGAGGA | | 7 | 458 | |
| MR54 | | CTGAGTC | | 7 | 475 | |
| MR55 | | TTTTGAAGGGAATTTTT | | 17 | 499 | 21 |
| MR56 | | ATTTTTA | | 7 | 510 | |
| MR57 | | TATGTAT | | 7 | 599 | |
| MR58 | | TAATAAT | | 7 | 605 | |
| MR59 | | TAGCATCGCTACTAT | | 15 | 670 | |
| MR60 | | ACTATCA | | 7 | 680 | |
| MR61 | | CAACCTACATCTAAC | | 15 | 788 | |
| MR62 | | TAAGAAT | | 7 | 836 | |
| MR63 | | AAATTATCTAAATCTCTTGAA | | 21 | 920 | |
| MR64 | | CATGGTAC | | 8 | 984 | |
| MR65 | | TACTCAT | | 7 | 989 | |
| REGION 4 | | | | | | |
| MR67 | | GCAAAACG | | 8 | 1 | |
| MR68 | | TATACAACATAT | | 12 | 60 | |
| MR69 | | GTTCTTG | | 7 | 82 | |
| MR70 | | ATTGAGCGAGCGATTTA | | 17 | 127 | |
| MR71 | | TTAATAATT | | 9 | 141 | |
| MR72 | | CTTTTTC | | 7 | 181 | |
| MR73 | | CGTTTGC | | 7 | 206 | |
| MR74 | | ATTATTA | | 7 | 238 | |
| MR75 | | AGATAGA | | 7 | 333 | |
| MR76 | | GCAAGTTTACAGCGCAA-ACTTGAACG | | 27 | 371 | |
| MR77 | | AGCGCAAACTTGAACGCGA | | 19 | 381 | |
| MR78 | | ATGAGTA | | 7 | 411 | |
| MR79 | | CCACACC | | 7 | 504 | 24 |
| MR80 | | TTAAAAATT | | 9 | 516 | |
| MR81 | | GAACAAG | | 7 | 527 | |
| MR82 | | TCAAAACT | | 8 | 594 | |
| MR83 | | AACTCTCAA | | 9 | 598 | |
| MR84 | | AATTTAA | | 7 | 609 | |
| MR85 | | TCCTTTCCT | | 9 | 621 | |
| MR86 | | GAGTGAG | | 7 | 715 | |
| MR87 | | TTAGTGATT | | 7 | 759 | |
| MR88 | | ACATTACA | | 8 | 805 | |
| MR89 | | GGTCTGG | | 7 | 918 | |
| MR90 | | GTCAACTG | | 8 | 928 | |
| REGION 5 | | | | | | |

| | | | | | | |
|---|---|---|---|---|---|---|
| MR91 | | TGCGTATGCGT | | 11 | 53 | |
| MR92 | | CTCTTCTC | | 8 | 81 | |
| MR93 | | TTCAACTT | | 8 | 157 | |
| MR94 | | CACAACAC | | 8 | 294 | |
| MR95 | | TCGAGCT | | 7 | 421 | |
| MR96 | | AACAGACAA | | 9 | 459 | |
| MR97 | | CAACCAAC | | 8 | 566 | 16 |
| MR98 | | TTGGGTT | | 7 | 620 | |
| MR99 | | TGAAAGT | | 7 | 626 | |
| MR100 | | GTTCTTG | | 7 | 639 | |
| MR101 | | GTGAGTG | | 7 | 679 | |
| MR102 | | GCTGGGTCG | | 9 | 711 | |
| MR103 | | AAGGGAA | | 7 | 769 | |
| MR104 | | TAGTCTGAT | | 9 | 864 | |
| MR105 | | TCGTTGCT | | 8 | 944 | |
| MR106 | | ATTCTTA | | 7 | 985 | |
| REGION 6 | | | | | | |
| MR107 | | ATACCATA | | 8 | 39 | |
| MR108 | | CGTTTGC | | 7 | 200 | |
| MR109 | | TACACAT | | 7 | 249 | |
| MR110 | | ATCCCAAGAACTCTA | | 13 | 276 | |
| MR111 | | GAGCGAG | | 7 | 318 | |
| MR112 | | TTCAATTGTTAACTT | | 15 | 377 | |
| MR113 | | ACTCTCA | | 7 | 399 | |
| MR114 | | AACATACAA | | 9 | 438 | |
| MR115 | | CAAAAAC | | 7 | 444 | |
| MR116 | | AATTTAA | | 7 | 513 | |
| MR117 | | AAGTTGAA | | 8 | 585 | |
| MR118 | | GAACAAG | | 7 | 602 | |
| MR119 | | ATATATA | | 7 | 669 | |
| MR120 | | CTGTGTC | | 7 | 700 | |
| MR121 | | AAGGGAA | | 7 | 712 | 26 |
| MR122 | | AAATAAA | | 7 | 719 | |
| MR123 | | GCCTTGAAGTTTCG | | 12 | 741 | |
| MR124 | | TGCTCGT | | 7 | 758 | |
| MR125 | | GGAAAAAGG | | 9 | 835 | |
| MR126 | | AAAAGGGAAAA | | 11 | 838 | |
| MR127 | | AAAGGGAAAAGGTAAA | | 14 | 839 | |
| MR128 | | GGTATGG | | 7 | 867 | |
| MR129 | | ATGGGTA | | 7 | 870 | |
| MR130 | | TACTCAT | | 7 | 924 | |
| MR131 | | CACTCAC | | 7 | 949 | |
| MR132 | | TAGAGAT | | 7 | 992 | |
| REGION 7 | | | | | | |
| MR133 | | AGAGAGA | | 7 | 4 | |
| MR134 | | AGAGATTTAGTGA | | 11 | 6 | |
| MR135 | | GTGAAGTG | | 8 | 15 | |
| MR136 | | AAAGAAA | | 7 | 25 | |
| MR137 | | ACAAAACA | | 8 | 165 | |
| MR138 | | AAACAAA | | 7 | 168 | |
| MR139 | | GAGAGAGAG | | 9 | 194 | |
| MR140 | | AGCTCGA | | 7 | 201 | |
| MR141 | | GAGGTGGAG | | 9 | 263 | |
| MR142 | | GCTAAATCG | | 9 | 278 | |
| MR143 | | AACCCAA | | 7 | 311 | |
| MR144 | | TGAAAGT | | 7 | 339 | |
| MR145 | | GGTTTTGG | | 8 | 377 | |
| MR146 | | ATACATA | | 7 | 394 | |
| MR147 | | ACCACCA | | 7 | 408 | |
| MR148 | | ATTTATTTA | | 9 | 414 | |
| MR149 | | TGGAGGT | | 7 | 441 | 29 |
| MR150 | | AGTTTGA | | 7 | 488 | |
| MR151 | | ATTTGTTTA | | 9 | 602 | |
| MR152 | | AAGAGAA | | 7 | 627 | |
| MR153 | | TGCTTCGT | | 8 | 637 | |
| MR154 | | TCATCACAGAAACAAG-CACTACT | | 20 | 648 | |

| | | | | | |
|---|---|---|---|---|---|
| MR155 | | ACCAGTGGTGAGCA | 12 | 736 | |
| MR156 | | CACACAC | 7 | 795 | |
| MR157 | | TTGGGTT | 7 | 883 | |
| MR158 | | GACACAG | 7 | 899 | |
| MR159 | | GTGTTGTG | 8 | 905 | |
| MR160 | | TTCAAAACAACAAAGCTT | 16 | 953 | |
| MR161 | | TGATGTAGT | 9 | 991 | |
| REGION 8 | | | | | |
| MR162 | | GCTGTCG | 7 | 58 | |
| MR163 | | GACACTTCACAG | 12 | 116 | |
| MR164 | | ATGCGTA | 7 | 185 | |
| MR165 | | TAGAGAT | 7 | 210 | |
| MR166 | | ATGCGTA | 7 | 215 | |
| MR167 | | ATGAAGTA | 8 | 235 | |
| MR168 | | CATCTAC | 7 | 308 | 17 |
| MR169 | | GTCACTG | 7 | 349 | |
| MR170 | | CAAAAAGAAAGAC | 11 | 417 | |
| MR171 | | GTGTGTG | 7 | 619 | |
| MR172 | | TGTGTGT | 7 | 620 | |
| MR173 | | TGGTGGT | 7 | 699 | |
| MR174 | | AGTTTGA | 7 | 767 | |
| MR175 | | TTGATAGTT | 9 | 770 | |
| MR176 | | ACTAACTCCATACCT-AATCA | 19 | 780 | |
| MR177 | | TCAGAAGC-ACATACATGGAAGACT | 21 | 815 | |
| MR178 | | TAATAAT | 7 | 936 | |
| REGION 9 | | | | | |
| MR179 | | GAGAAAGAG | 9 | 96 | |
| MR180 | | AGAACAAGA | 9 | 168 | |
| MR181 | | AAGAAGGAAGGAGGAA | 14 | 173 | |
| MR182 | | GCTAATCG | 8 | 215 | |
| MR183 | | AAGAGAGAA | 9 | 258 | |
| MR184 | | AGATTAGA | 8 | 303 | |
| MR185 | | GGAAGGGAAGG | 11 | 422 | |
| MR186 | | GAGTTTGAG | 9 | 537 | |
| MR187 | | ATGAAGTA | 8 | 556 | |
| MR188 | | AAAGGAAA | 8 | 654 | |
| MR189 | | AGGAAAGGA | 9 | 656 | |
| MR190 | | CACAGTTTGATAC | 11 | 835 | 25 |
| MR191 | | TGGTTTGGT | 9 | 919 | |
| MR192 | | TGAACAAGT | 9 | 986 | |
| MR193 | | CGTATATAGAAATGCGCAACAAAAAGGAACCATATATGC | 27 | 1078 | |
| MR194 | | ACGGTGGCA | 9 | 1279 | |
| MR195 | | ACACAAGAGGAGAACACA | 18 | 1293 | |
| MR196 | | TGTGATGTAGTGT | 13 | 1372 | |
| MR197 | | ATAATTAATA | 10 | 1454 | |
| MR198 | | CTTAGCTGTCGGTTC | 13 | 1523 | |
| MR199 | | TTCTGTATTATTAAGTCTT | 17 | 1535 | |
| MR200 | | GTTGTTGTTG | 10 | 1570 | |
| MR201 | | TGTTGTTGT | 9 | 1572 | |
| MR202 | | TAGCAGTGACTAT | 11 | 1621 | |
| MR203 | | AACTTCAA | 8 | 1688 | |
